# Divergent Protein Kinase A contributes to the regulation of flagellar waveforms in *Leishmania mexicana*

**DOI:** 10.1101/2023.11.08.566240

**Authors:** Sophia Fochler, Benjamin J Walker, Richard J Wheeler, Eva Gluenz

## Abstract

The second messenger cyclic adenosine monophosphate (cAMP) and protein kinase A (PKA) have been implicated in the regulation of flagellar motility in diverse eukaryotes, yet the specific contributions of PKA isoforms remain unclear. Moreover, the kinetoplastid PKA is insensitive to cAMP. Here we investigated subcellular localisations of PKA subunits in *Leishmania* and study the flagellar waveform parameters in PKA deletion mutants. Expansion microscopy localised *Lmx*PKAR1 between the paraflagellar rod and the axoneme and *Lmx*PKAC1, *Lmx*PKAC2 depend on *Lmx*PKAR1 for enrichment in the flagellum. *Lmx*PKAC3 requires *Lmx*PKAR3 for co-localisation at the cell cortex. Deletion of *Lmx*PKAR3 resulted in increased flagellar *Lmx*PKAC3 signal and increased swimming speed. Conversely, deletion of *Lmx*PKAC3 resulted in reduced flagellar beat frequencies and swimming speed. The absence of *Lmx*PKAC1 resulted in a lower incidence of high frequency tip-to-base waveforms, and consequently reduced swimming speed. *Lmx*PKAC1 dissociated from cytoskeletons upon addition of guanosine, adenosine or inosine, consistent with a novel nucleoside-based PKA activation mechanism. These data suggest that this divergent PKA pathway regulates the motility of *Leishmania* through the modulation of flagellar waveforms.

## Introduction

Motile cilia and flagella are dynamic, specialised organelles used by many cells for movement through the environment. Over 700 proteins (Broadhead et al. 2006; Nakachi et al. 2011; Pazour et al. 2005; Vasquez, van Dam, and Wheway 2021) form the flagellar cytoskeleton, the axoneme, which typically consists of nine doublet microtubules and two central singlet microtubules (called the central pair, CP). Radial spoke complexes (RS) project from the doublets to the CP and the nexin links/dynein regulatory complex (N-DRC) project from one doublet to its neighbour radially. Two rows of dynein motor protein complexes, the outer dynein arms (ODA) and inner dynein arms (IDA), align in a 96 nm repeating pattern along the doublet microtubules (Lindemann and Lesich 2010). Dyneins are a family of ATPases which hydrolyse ATP to iteratively bind, pull and release adjacent microtubules (Heale and Alisaraie 2020). This sliding is converted to bending by sliding restrictions imposed by the basal body and sliding resistance from the N-DRC (Heuser et al. 2009). While eukaryotic flagella have the same conserved core structures, important open questions remain about how flagellar waveforms and frequency are modulated. Moreover, little is known of how different cells translate cues from the local environment into appropriate flagellar waveforms.

Calcium and cAMP are the two main intracellular signalling molecules (called second messengers) that have been linked to changes in flagellar waveforms in diverse types of cilia and flagella, including mammalian respiratory cilia (Nlend et al. 2007), human sperm (Chung et al. 2017; Balbach et al. 2018), bull sperm (Alonso et al. 2017; Ishijima 2015), hamster sperm (Aoki, Sakai, and Kohmoto 1999; Kinukawa et al. 2006) and sea urchin sperm (Tash 1989; Vacquier et al. 2014; Beltrán, Zapata, and Darszon 1996), the green algae *Chlamydomonas* (Hyams and Borisy 1978; Saegusa and Yoshimura 2015; Schmidt and Eckert 1976), *Paramecium* (Lodh et al. 2016; N M Bonini, D L Nelson 1988), Euglena (Nasir et al. 2022) and *Leishmania* (Mukhopadhyay and Dey 2016; 2017). Intraciliary cAMP concentration correlates with beat frequency in mammalian cilia (Schmid et al. 2006; Ma, Silberberg, and Priel 2002). The second messenger signal is transmitted through kinases, phosphatases and calcium-binding proteins that are have been found to be incorporated into core axoneme structures (Tash 1989; Christensen et al. 2001; Brokaw and Kamiya 1987; Kutomi et al. 2012; Hamasaki et al. 1991). A key transducer of cAMP signals is cAMP-dependent protein kinase (PKA), a highly conserved protein sharing similarities with cGMP-dependent protein kinase (PKG) and protein kinase C (PKC) (Vacquier et al. 2014), collectively named the AGC kinase subgroup. The paradigm for eukaryotic PKA signal transduction involves a heterotetrameric holoenzyme composed of two PKA catalytic subunits (PKAC) and two PKA regulatory subunits (PKAR), (C_2_R_2_). In vertebrates, A-kinase anchoring proteins (AKAPs) bind the PKA heterotetrameric holoenzyme via the RII domains in the PKAR, creating local micro-signalosomes (Torres-Quesada, Mayrhofer, and Stefan 2017). However, most recently, mammalian PKA has been shown to also exist in homodimer form (C_1_R_1_) localising to radial spoke 3 within bovine sperm axonemes (Leung et al. 2025). In its inactive state, unbound to cAMP, the autoinhibitory domain (AI) of PKAR occupies and inhibits the catalytic core of the PKAC subunit (Anand et al. 2007). Upon binding of a cAMP molecule to each of the PKAR subunits, the PKAC is then released to phosphorylate its substrates. Flagellar ODAs are thought to represent an important target of PKA (Peng et al. 2015a; Salathe 2007).

Flagellate protozoa of the genera *Leishmania* and *Trypanosoma* are well-studied due to their importance as parasites of humans and livestock. They also provide genetically tractable study models for eukaryotic flagella in a lineage that is distinct from the three lineages in which humans, green algae (*Chlamydomonas*) or ciliates (e.g. *Tetrahymena*, *Paramaecium*) evolved. Motile *Leishmania* use a symmetric tip-to base flagellar waveform to move forward, and switch to an asymmetric base-to-tip waveform to rotate the cell and change direction (Gadelha, Wickstead, and Gull 2007). Trypanosomatid signal transduction pathways remain poorly defined and, whilst cAMP causes flagellar waveform switching in *Leishmania* (Mukhopadhyay and Dey 2016), there is considerable uncertainty in how signals are transduced within trypanosomatid flagella.

All kinetoplastid genomes available on VEuPathDB (Alvarez-Jarreta et al. 2024) asides from the aflagellate endosymbiont *Perkinsela*, encode three PKAC genes that share some canonical features of eukaryotic PKAs, such as the conserved phosphoresidue located in the activation loop (Cheng, Zhang, and McCammon 2006) along with a conserved catalytic core sequence (Bachmaier et al. 2019a). Intriguingly, PKA in trypanosomes forms C_1_R_1_ heterodimers and does not respond to cAMP (Bachmaier et al. 2019a; Shalaby, Liniger, and Seebeck 2001). Instead, nucleosides and nucleoside analogues, including inosine, 7-deazaadenosine (tubercidin) and toyocamycin, bind trypanosomatid PKAR with high affinity (Ober et al. 2024). Interestingly, when the known mammalian PKA phosphorylation substrate VASP was exogenously expressed in *T. brucei,* it was phosphorylated on serine-157 by endogenous parasite PKA (Bachmaier et al. 2019a), suggesting that, although *Trypanosoma* PKA homologues are not regulated by cyclic nucleotides, their target site specificity may still be shared across eukaryotic species.

Several lines of evidence point towards the trypanosomatid flagellum as a key site for PKA function. First, flagellar localisation has been demonstrated for PKAC1/2 and PKAR subunits, in flagellar proteomes of *T. brucei* and *L. mexicana* (Oberholzer et al. 2011; Beneke et al. 2019), by labelling *L. donovani* promastigote forms and *T. brucei* with PKAR1-specific antibodies (Bachmaier et al. 2016) and in protein tagging screens (Beneke et al. 2019; Dean, Sunter, and Wheeler 2017; Billington et al. 2023). Moreover, the flagellar localisation of TbPKAR1 was shown to depend on IFT (Fort et al. 2016). Second, loss-of-function mutations altered parasite motility patterns: depletion of TbPKAR1 by RNAi in bloodstream forms reduced the cells’ capacity for forward motility (Oberholzer et al. 2011). A complete deletion of PKAR1 or a partial deletion of PKAC1 slowed the swimming speeds of *L. mexicana* promastigotes (Beneke et al. 2019). Finally, the developmental regulation of transcripts and activity in *Leishmania* spp. reflects the presence or absence of a motile flagellum: *L. major* promastigotes, which possess a motile flagellum, showed much higher transcript levels (Siman-Tov et al. 1996) than the amastigote form, which only have a short, non-motile cilium (Bachmaier et al. 2016; Kobe and Kajava 2001). Proteome-wide, the phosphorylation kinetics of the PKA target motif (RXXS*/T*) in *L. donovani* during promastigote to amastigote differentiation consistently detected the highest signal in promastigote forms (Bachmaier et al. 2016), suggesting a promastigote-specific function for PKA activity. Still, the precise PKA localisation, the mechanism of upstream physiological activation, and downstream biological function remain to be fully explored.

Here we present a systematic analysis of three PKAC and two PKAR isoforms of *L. mexicana*, using gene deletions and microscopy to discover how gene loss affects flagellar beat dynamics. Localisation of fluorescently tagged proteins showed distinct subcellular localisation patterns for PKAC and PKAR subunits, and specific co-dependencies, from which we infer pairing preferences for *Lmx*PKAC1 and *Lmx*PKAC2 with *Lmx*PKAR1 in the flagellum, and *Lmx*PKAC3 with *Lmx*PKAR3 along the cell cortex. Ultrastructure expansion microscopy showed that LmxPKAR1 is localised adjacent to the axonemal structure, within the inner/middle domain of the paraflagellar rod. Targeted gene deletion of PKA subunits individually and in combination showed that none are essential for promastigote growth. Assessment of swimming speed and measurement of flagellar waveform parameters showed that LmxPKAR3/C3 seem to modulate beat frequency and swimming speed, whereas *Lmx*PKAC1 is essential for the generation of a symmetric tip-to base flagellar waveform.

## Results

### Leishmania spp. have three PKAC and two PKAR subunits

BLASTp searches with the three PKAC and the single PKAR protein sequences from *Trypanosoma brucei* (Bachmaier et al. 2019a) identified syntenic *Leishmania mexicana* orthologues for all four encoding genes (Table 1, Supplementary Figure 1A-C) (Amos et al. 2022). Analysis of the *L. mexicana* PKAC sequences confirmed overall conservation of the trypanosomatid PKAC repertoire: the highly similar *Lmx*PKAC1 and *Lmx*PKAC2 are located on chromosome 34, not in a tandem array but separated by four other genes. This locus is syntenic with the locus of *T. brucei* and *T. cruzi* PKAC1 and PKAC2, but the designation of these PKACs as 1 or 2, respectively, varies between species (Table 1, Supplementary Figure 1A). *Lmx*PKAC1 and *Lmx*PKAC2, but not *Lmx*PKAC3, have variable N-terminal extensions that harbour a predicted disordered domain (Supplementary Figure 2A, B) and both the active sites and peptide binding sites within all *Leishmania* PKAC1 and PKAC2 are identical (Supplementary Figure 2C, D). Catalytic activity of eukaryotic PKAC is modulated by phosphorylation of several residues, such as threonine-197 in the activation loop, which is particularly involved in regulating activity (Steinberg et al. 1993; Steichen et al. 2012), and is conserved across trypanosomatid PKAC (Supplementary Figure 2E). At the C-terminus, trypanosomatid PKAC1 and PKAC3 have a conserved FXXF sequence, where mutating either phenylalanine was shown to diminish catalytic activity (Etchebehere et al. 1997; Chan et al. 2020). *LmjF*PKAC2 lacks the first of these phenylalanines (Supplementary Figure 2F). All trypanosomatid PKAC3, including *Lmx*PKAC3, have an 8 amino acid C-terminal extension of unknown function (Siman-Tov et al. 1996) (Supplementary Figure 2F).

**Table 1.**
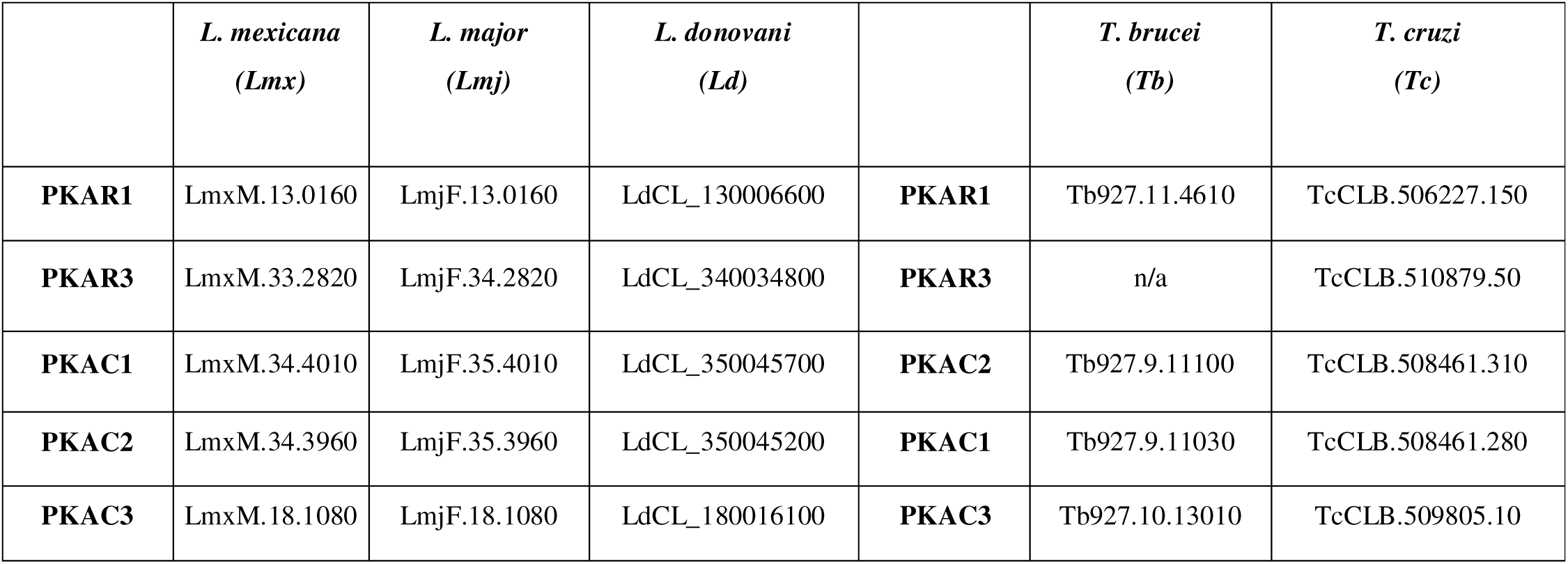
Accession numbers for all PKAR and PKAC orthologues found encoded in the genomes of *L. mexicana, L. major, L. donovani, T. brucei* and *T. cruzi*. Each row represents genes found in a syntenic locus across kinetoplastid species, as illustrated in Supplementary Figure 1A-D.

|  | *L. mexicana*<br>(*Lmx*) | *L. major*<br>(*Lmj*) | *L. donovani*<br>(*Ld*) |  | *T. brucei*<br>(*Tb*) | *T. cruzi*<br>(*Tc*) |
| --- | --- | --- | --- | --- | --- | --- |
| **PKAR1** | LmxM.13.0160 | LmjF.13.0160 | LdCL_130006600 | **PKAR1** | Tb927.11.4610 | TcCLB.506227.150 |
| **PKAR3** | LmxM.33.2820 | LmjF.34.2820 | LdCL_340034800 | **PKAR3** | n/a | TcCLB.510879.50 |
| **PKAC1** | LmxM.34.4010 | LmjF.35.4010 | LdCL_350045700 | **PKAC2** | Tb927.9.11100 | TcCLB.508461.310 |
| **PKAC2** | LmxM.34.3960 | LmjF.35.3960 | LdCL_350045200 | **PKAC1** | Tb927.9.11030 | TcCLB.508461.280 |
| **PKAC3** | LmxM.18.1080 | LmjF.18.1080 | LdCL_180016100 | **PKAC3** | Tb927.10.13010 | TcCLB.509805.10 |

Several reports have highlighted the existence of a second PKAR-like protein within the trypanosomatids (Baker et al. 2021; Jäger et al. 2014), which was given the name PKAR3 in *L. donovani* (Fischer Weinberger et al. 2024); for consistency, we adopt this terminology here*. L. mexicana* has a syntenic orthologue of PKAR3 (Table 1) (Supplementary Figure 1D). All trypanosomatid PKAR3s share a similar protein architecture with canonical PKARs (Figure 1A), with two CNBDs towards the C-terminus and an autoinhibitory (AI) domain that mimics the phosphorylation targets of PKAC and inhibits the catalytic kinase subunits. This critical sequence is typically RRXA(G) in Type I PKARs (PKAR-I) and RRXS(T) in Type II PKARs (PKAR-II) (Figure 1B). In Type I PKARs, the alanine (A) or Glycine (G) replaces the phosphorylatable serine (S) or threonine (T) residues found in substrate sites (C. E. Poteet-Smith et al. 1997; Scott et al. 1985). Trypanosomatid PKAR1 sequences all have an RRXT sequence that aligns with the autophosphorylatable RRXS motif in human PKAR-II. The *Leishmania* PKAR3 sequences share the RGAIS motif of the canonical mammalian RRRRGAISAE sequence of the PKAR-Iα AI, but notably are missing the first arginine residue of the Type I PKAR motif RRXA (Figure 1B).

**Figure 1.**
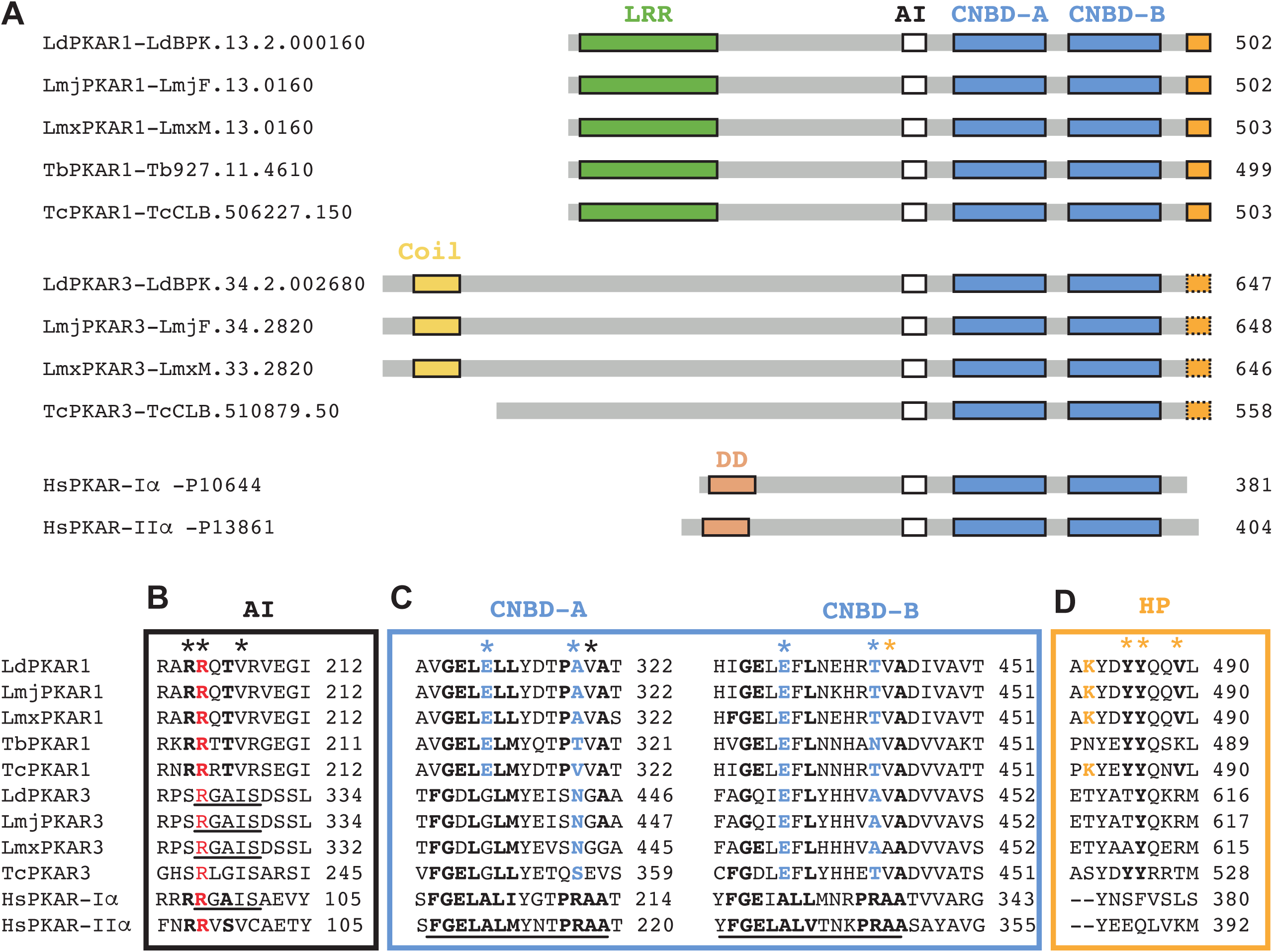
Alignments of PKAR sequences. **(A)** Cartoon representation of trypanosomatid PKAR proteins *Ld*PKAR1 (LdBPK.13.2.000160), *Lmj*PKAR1 (LmjF.13.0160), *Lmx*PKAR1 (LmxM.13.0160), *Tb*PKAR1 (Tb927.11.4610), *Tc*PKAR1 (TcCLB.506227.150), *Ld*PKAR3 (LdBPK.34.2.002680), *Lmj*PKAR3 (LmjF.34.2820), *Lmx*PKAR3 (LmxM.33.2820), *Tc*PKAR3 (TcCLB.510879.50), and Homo sapiens cAMP-dependent protein kinase type I-alpha regulatory subunit PKAR-Iα (P10644, KAP0_HUMAN) and type II-alpha regulatory subunit PKAR-IIα (P13861|KAP2_HUMAN). LRR, leucine rich repeat; DD, dimerization docking domain; CNBD, cyclic nucleotide binding domains (PFAM ID: PF00027); AI, autoinhibitory domain sequence (Celeste E. Poteet-Smith et al. 1997). **(B)** AI sequence alignment. Bold letters highlight the RRxA and RRxT motifs which form a pseudo-substrate or are autophosphorylated by docked catalytic subunits in mammalian PKA heterotetramers. A universally conserved arginine residue is highlighted in red. The conserved RGAIS residues in *Leishmania* PKAR3 proteins and the human AI sequence are underlined. **(C)** CNBD: Underlined is the canonical phosphate binding cassette (PBC) motif FGELAL[ILVM]xxxPRAA (Peng et al. 2015b); residues matching the PBC are in bold black type. Highlighted in blue (asterisks) are residues associated with altered ligand specificity in trypanosomatid PKAR1 (Bachmaier et al. 2019b): the switch from alanine (position 5) in human PKA to glutamate in trypanosomatid and the switch from arginine (position 12) in human PKA to a alanine, valine, serine, threonine, or asparagine in the trypanosomatid PKAR sequences. Orange asterisk indicate a valine residue reported to form part of the trypanosomatid specific hydrophobic pocket. **(D)** Hydrophobic pocket: orange asterisk indicates a residue reported to form part of the *Trypanosoma* specific hydrophobic pocket. Highlighted in orange type is a lysine residue reported to increase affinity to binding synthetic nucleoside analogue 7-cyano-7-deazainosine (Bachmaier et al. 2019b).

Deviations from the canonical PKAR architecture are found at the N-termini (Figure 1A): trypanosomatid PKAR1 proteins lack the dimerization/docking domain (DD; PF02197) that holds canonical eukaryotic PKA heterotetramer together and anchors it to the amphipathic helix region of an A-kinase anchoring protein (AKAP) (Banky, Huang, and Taylor 1998). Instead, they contain a predicted leucine-rich repeat (LRR; IPR032675). PKAR3 sequences lack a recognisable DD domain or an LRR and, instead, there is a short coil domain predicted near the N-terminus of all examined PKAR3 sequences. This suggests that the family of divergent kinetoplastid PKAR have evolved subcellular anchoring mechanisms distinct from AKAP.

### The cyclic nucleotide binding domains of Leishmania PKAR subunits deviate from the canonical cAMP binding domain of mammalian PKAR

Studies using crystal structures showed that, in all trypanosomatid PKAR1s, there are substitutions to key residues in the phosphate binding cassette (PBC) that impede docking of the cAMP molecule (Bachmaier et al. 2019a): a conserved alanine (A) is replaced by a negatively charged glutamate (E), which is predicted to clash with the phosphate group on the cAMP molecule, and the arginine (R) residue required to dock the exocyclic phosphate has been switched to polar uncharged (R/TN) or aliphatic (R/AV) residues (Bachmaier et al. 2019a; Ober et al. 2024). These changes have been experimentally demonstrated to cause a ligand switch from cAMP to nucleoside analogues in trypanosomatid PKAR1s, and they are conserved in *Leishmania* PKAR1 sequences (Figure 1C). These findings prompted us to examine in detail the nucleotide binding sites of the PKAR3 subunits. Key residue switches are evident in the parasite PKAR3 CNBDs but with some notable deviations from the PKAR1 patterns (Figure 1C). In the place of the first alanine (A) in CNBD-A, PKAR3 has a glycine (G), which, assuming a local neutral pH environment would have a net zero charge. Thus, it is hard to predict if this residue would clash with cAMP docking in the same manner as the glutamate in PKAR1, defined by (Bachmaier et al. 2019a). The first alanine in CNBD-B has been substituted for a glutamate in PKAR3, identical to PKAR1 sequences, and the arginine residues have also been substituted in PKAR3 CNBD-A and CNBD-B with polar uncharged (R/STN) or hydrophobic side chains (R/A). Most surprisingly is that the conserved initial glutamate within the PBC (G**<u>E</u>**LAL), which forms a hydrogen bond with the 2’-OH moiety on the ribose sugar of cAMP (Berman et al. 2005), has been switched to an aspartic acid (D) in the *Leishmania* PKAR3 CNBD-A and aspartic acid or glutamine (D/Q) in all PKAR3 CNBD-B asides from *Lmx*PKAR3. These changes suggest that PKAR3, like PKAR1, has diverged from cAMP binding, but raise the question of whether the same or different ligands bind to each of these sites.

All parasite sequences for PKAR1 and PKAR3 have an extended C-terminus end compared to the canonical mammalian PKAR reference sequence (Figure 1A). Two tyrosine (Y) and two valine (V) residues were implicated in forming a parasite specific hydrophobic pocket documented in *Tc*PKAR1 (V444, Y486, Y485 and V489 denoted by orange asterisks in Figure 1C and D), interactions with which were shown to increase the binding affinity and activation potential of certain nucleoside analogues (Bachmaier et al. 2019a; Ober et al. 2024). All parasite PKAR1 sequences, contain at least two of these YV residues. Contrastingly, in PKAR3 sequences, *Leishmania* only have one Y and one V, where *Tc*PKAR3 has three of the four hydrophobic residues. Taken together these data indicate that a divergent PKAR1 is conserved in all trypanosomatids, and an additional PKAR3 is found in *Leishmania* spp. and *T. cruzi*, with an unknown ligand specificity, that shares in its CNBD sequences some of the divergent features of PKAR1.

### L. mexicana PKACs and PKARs show distinct localisation patterns to the flagellum and cell body cortex

To establish the subcellular localisation of the different PKAC and PKAR subunits in *L. mexicana* in different cell- and life-cycle stages, we inserted fluorescent protein fusion tags (mNeonGreen, mNG) at the endogenous gene loci, producing mutants expressing either N- or C-terminal fusion products for each PKA subunit.

mNG::PKAC1 localised to the flagellum in non-dividing cells (1 kinetoplast, 1 nucleus, 1 flagellum (1K1N1F) (Wheeler, Gluenz, and Gull 2011), a signal that remained strong in 0.1% NP-40 detergent-extracted cytoskeletons (Figure 2A). Through the cell division cycle, the signal abundance diminished in the flagellum and increased within the nucleus (Supplementary Figure 3A), corroborating the nuclear localisation that was reported for this protein in a *L. mexicana* kinase localisation screen (Baker et al. 2021). Strikingly, within the nucleus, the *Lmx*PKAC1 signal intensity was consistently strongest in areas of low Hoechst staining (Supplementary Figure 3A’), indicating localisation to the nucleolus. The *Lmx*PKAC1 signal re-appeared in both the old and the newly growing flagella at cytokinesis. In axenic amastigotes, the mNG::PKAC1 signal appeared greatly diminished, with weak variable cytoplasmic signal (Figure 2A). A C-terminal tag (PKAC1::mNG) disrupted the localisation to the flagellum (Supplementary Figure 4A), as previously reported (Beneke et al. 2019).

**Figure 2.**
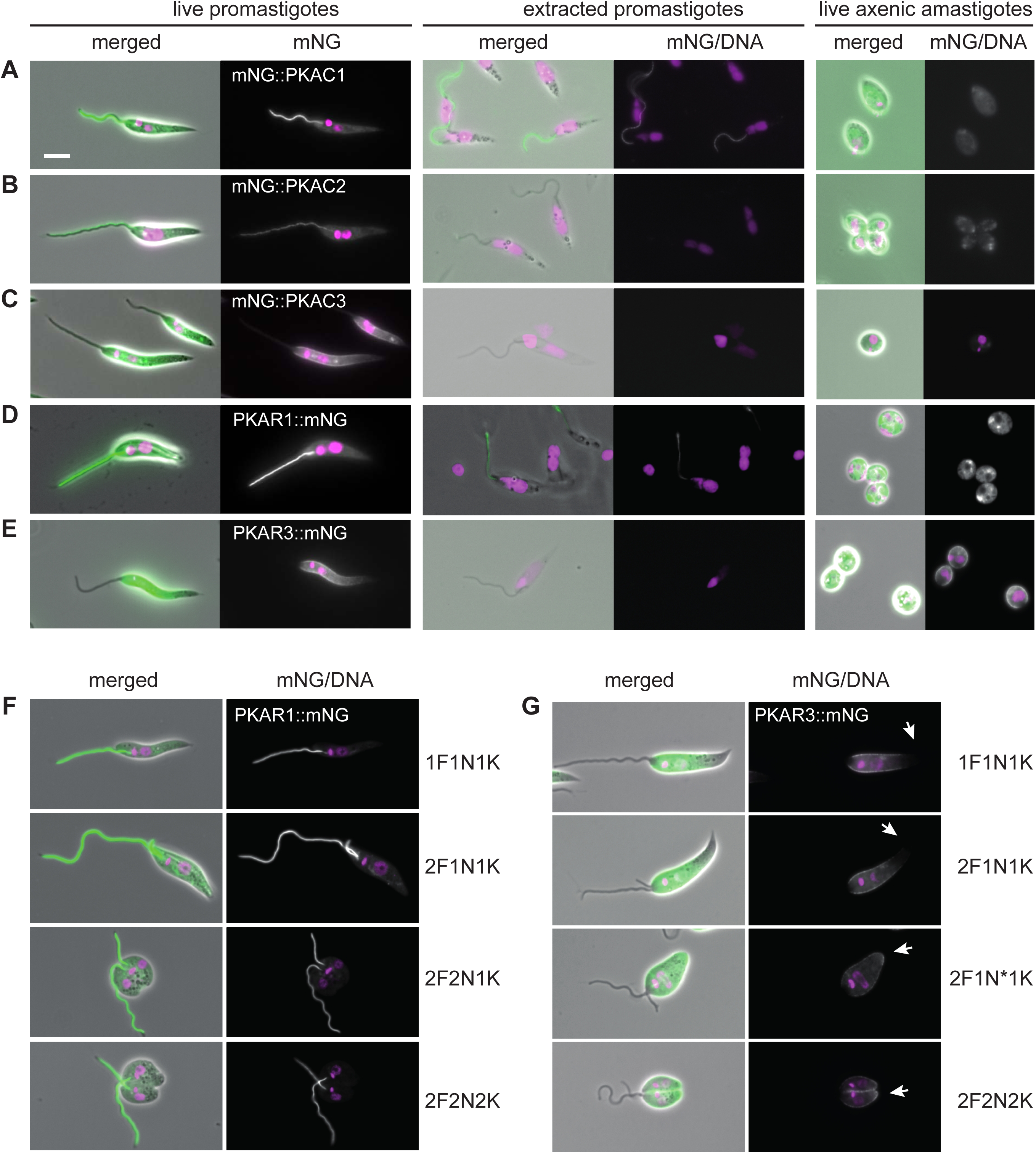
Localisations of tagged *L. mexicana* PKAC and PKAR subunits. Micrographs of cells expressing mNG-tagged PKAC or PKAR subunits. **(A)** mNG::*Lmx*PKAC1 (LmxM.34.4010), **(B)** mNG::*Lmx*PKAC2 (LmxM.34.3960), **(C)** mNG:: *Lmx*PKAC3 (LmxM.18.1080), **(D)** *Lmx*PKAR1::mNG (LmxM.13.0160), **(E)** *Lmx*PKAR3::mNG (LmxM.33.2820). Panels in A-E show live promastigotes (left), detergent-extracted promastigote cytoskeletons (middle) and live axenic amastigotes (right). For each sample type a merged fluorescence with phase contrast image is shown, and the combined mNG and Hoechst fluorescence. **(F)** Promastigotes in different stages of the cell cycle, expressing *Lmx*PKAR1::mNG and **(G)** mNG::*Lmx*PKAR3. White arrows indicate dynamic localization of mNG::*Lmx*PKAR3 to posterior cell body pole. F, flagella; N, nuclei; N*, dividing nucleus, K, kinetoplasts. Scale bar indicates 5 µm.

The mNG::PKAC2 signal was partitioned between the flagellum, the cell body and cortex throughout in the promastigote, with variations in relative intensity in the different compartments at different cell cycle stages (Figure 2B and Supplementary Figure 3B). Notably, in early promastigote division (2F1N1K), the *Lmx*PKAC2 signal appeared enriched in the new growing flagellum and a strong cell cortex signal was seen during mitosis (Supplementary Figure 3B). The *Lmx*PKAC2 axonemal signal was lost from detergent-extracted cytoskeletons, and only a weak variable cytoplasmic signal was seen in axenic amastigotes (Figure 2B). Fusion of mNG to the C-terminus hindered protein localisation to the flagellum (Supplementary Figure 4B).

The mNG::PKAC3 signal was associated with the cell body cortex in non-dividing cells, with a signal that was strongest at the central region of the cell body and tapering off towards the posterior pole (Figure 2C). In 95% of non-dividing cells, a distinctive ‘edge effect’ was evident (Figures 2C and 3A) (Halliday et al. 2019). A faint signal was also present in the flagellum, with an intensity gradient decreasing towards the distal flagellum (Figure 2C). This signal was lost upon membrane extraction with 0.1% NP-40 (Figure 2C, middle panel). In dividing promastigotes, the *Lmx*PKAC3 signal appeared more uniform across the cell body cortex, with an accumulation at the anterior end of the developing cleavage furrow and a reduced flagellar signal (Supplementary Figure 3C). The promastigote cortex signal disappeared in detergent extracted cells and showed a weak variable cytoplasmic signal in the cell body of axenic amastigotes (Figure 2C).

**Figure 3.**
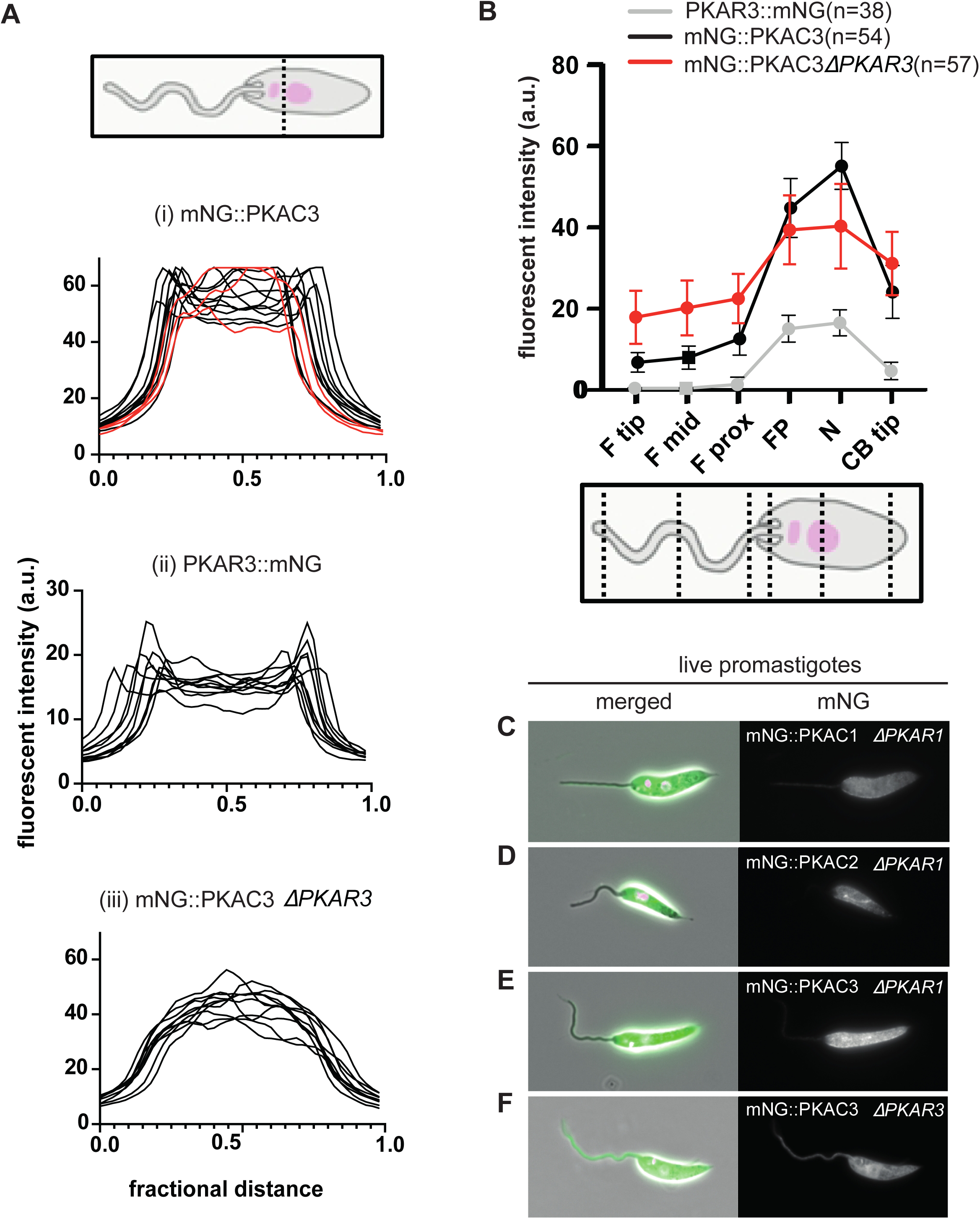
Localisation dependencies of *L. mexicana* PKAC and PKAR subunits. **(A)** Measurement of the mNG fluorescent signal intensity across a line though the cell body (cartoon) show a clear edge effect for the majority of mNG::*Lmx*PKAC3 cells (i) (black lines, n = 11); the red lines show a minority of cells (n = 3) with a diffuse cell body signal. An edge effect was detected for all measured *Lmx*PKAR3::mNG cells (ii) (n = 12). The edge effect for mNG::*Lmx*PKAC3 was lost following *Lmx*PKAR3 deletion (iii) (n = 10). **(B)** Quantitation of fluorescence signal intensity in cells expressing *Lmx*PKAR3::mNG, mNG::*Lmx*PKAC3 and in cells expressing mNG::*Lmx*PKAC3 following *Lmx*PKAR3 deletion. Measurements were taken at six points along the flagellum and cell body as shown in the cartoon. **(C-F)** Micrographs showing the localisation of mNG-tagged *Lmx*PKAC subunits following deletion of *Lmx*PKAR subunits: **(C)** mNG::*Lmx*PKAC1 △*LmxPKAR1*; **(D)** mNG::*Lmx*PKAC2 △*LmxPKAR1*; **(E)** mNG::*Lmx*PKAC3 △*LmxPKAR1*; **(F)** mNG::*Lmx*PKAC3 △*LmxPKAR3*.

PKAR1::mNG showed a strong flagellar signal in promastigote forms, with a fainter diffuse cell body signal throughout all stages of promastigote cell division (Figure 2D, 2F). Upon detergent extraction, the fluorescent signal remained in the axonemal cytoskeleton. Fusion of mNG to the N-terminus of PKAR1 resulted in a weak reticulated cell body signal, suggesting that blocking the LRR domain may affect protein targeting (Supplementary Figure 4D). After induction of axenic amastigote differentiation for three days, a strong PKAR1::mNG signal was observed in the cytoplasm of axenic amastigotes (Figure 2D).

Both N- and C-terminally tagged *Lmx*PKAR3 localised exclusively to the cell body cortex in live promastigotes (Figure 2E and Supplementary Figure 4E). This signal was detergent extractable (Figure 2E), suggesting *Lmx*PKAR3 is anchored weakly to the cortical cytoskeleton, or is associated with the plasma membrane. In axenic amastigotes, the *Lmx*PKAR3 signal persisted at the cell body cortex (Figure 2E). Closer examination of the different cell cycle stages showed that, in non-dividing (1F1N1K), and early dividing (2F1N1K) promastigote cells, the cortical *Lmx*PKAR3 signal was most intense in the cell body mid-zone and tapered off towards the posterior pole (Figure 2G, quantified in Figure 3B). In late division (2F2K1N / 2F2K2N), the *Lmx*PKAR3 signal intensity became more uniform along the cortex, occasionally appearing enriched at the posterior pole (Figure 2G).

Overall, these data show that PKA concentrations in *L. mexicana* vary across sub-cellular space and over time, with two dominant PKA structures being the promastigote flagellum (dominated by *Lmx*PKAC1, -C2 and -R1) and the cell cortex of promastigotes and amastigotes (*Lmx*PKAC3 and -R3). Within these dominant structures, there were cell-cycle stage dependent changes in signal intensity and also a transient shift of the *Lmx*PKAC1 signal in dividing cells into the nucleus and away from the flagellum.

### Identification of localisation dependencies between PKAR and PKAC subunits to the flagellum and cell cortex

Given the striking similarities in the localisation of *Lmx*PKAC3 and *Lmx*PKAR3 at the cell cortex (Figure 2C, 2E), and the strong flagellar localisation of *Lmx*PKAC1 and *Lmx*PKAR1, the question arose whether there were specific PKAC-PKAR pairings in different cellular compartments. To test whether the localisation of any PKAC subunit was dependent on the expression of specific PKARs, we deleted the *Lmx*PKAR1 or *Lmx*PKAR3 genes in cells expressing tagged *Lmx*PKAC1, *Lmx*PKAC2 or *Lmx*PKAC3. Following deletion of *Lmx*PKAR1, promastigotes no longer showed the strong axonemal *Lmx*PKAC1 signal that was seen in the parental cell line (Figure 3C). Similarly, the flagellar signal from *Lmx*PKAC2 was also diminished upon deletion of *Lmx*PKAR1 (Figure 3D). By contrast, deletion of *Lmx*PKAR1 had no visible effect on *Lmx*PKAC3 localisation at the cell cortex (Figure 3E). We then tested whether loss of *Lmx*PKAR3 would affect *Lmx*PKAC3 localisation and found that the distinct cortical signal of *Lmx*PKAC3 became diminished and the signal appeared to become more widely distributed throughout the cell body (Figure 3F). This change in signal distribution was quantified over a cross-section of the cell body between the kinetoplast and the nucleus. This showed a complete loss of the edge effect for *Lmx*PKAC3 signal after *Lmx*PKAR3 was deleted (Figure 3A, iii). Secondly, the change in signal distribution along the axis of the cell was assessed by measuring fluorescent intensity at six positions along the cell (Figure 3B). In a parental background, the signal profile of *Lmx*PKAC3 was at its lowest at the distal flagellum, gradually increasing in intensity towards the pocket region (Figure 3B). Following deletion of *Lmx*PKAR3, the *Lmx*PKAC3 signal increased in the flagellum and decreased in the cell body (Figure 3B). Taken together, these data suggest pairing preferences between PKAC and PKAR subunits, whereby in non-dividing promastigotes, *Lmx*PKAR1 anchors *Lmx*PKAC1 and *Lmx*PKAC2 in the flagellum, whilst *Lmx*PKAR3 docks *Lmx*PKAC3 at the cell body cortex.

### PKAR1 is located in the zone between the axoneme and paraflagellar rod

We next tested whether the *Lmx*PKAR1 was located at the radial spokes of the *Leishmania* axoneme, since it was suggested that PKA associated with the AKAP domain of RSP3 in *Chlamydomonas* spokes 1 and 2 (Gaillard et al. 2001; Yang and Yang 2006; Gaillard et al. 2006). More recently, a cryo-EM structure directly showed mammalian PKA associated with malate dehydrogenase in the head of radial spoke 3 in human cilia (Leung et al. 2025). We deleted proteins of the head (RSP9) and stalk (RSP3 and RSP11) of *L. mexicana* radial spokes 1 and 2 (Dahlin, Zheng, and Scott 2021) in the PKAR1::mNG expressing cell line to see if the flagellar signal disappeared. A PKAR1::mNG signal was still observed in the axoneme of the examined RSP deletion mutants, including the short flagella that are characteristic of RSP3 deletion mutants (Supplementary Figure 5A-D). We next employed ultrastructure expansion microscopy to visualise the PKAR1 localisation patterns within *Leishmania* flagella (Figure 4) at higher resolution. Cells were stained with antibodies against the c-myc epitopes (contained in the PKAR1::mNG fusion protein), alpha-tubulin and PFR2 (a marker for the outer domain of the PFR (Kohl, Sherwin, and Gull 1999)). In the expanded cells, microtubules of the axonemes were clearly resolved and the PFR2 signal was on one side of the axoneme, adjacent to the tubulin signal. The PKAR1 signal was sandwiched between these two reference structures (Figure 4A, 4B). A recent systematic study of paraflagellar rod (PFR) sub-domains defined in *T. brucei* predicted PKAC1, PKAC2 and PKAR1 to reside within the inner/middle domain of the PFR (Gabriel et al. 2024). The observed fluorescence pattern is consistent with this prediction. The PKAR1 signal initiated in the flagellar pocket, before that of PFR2, and extended beyond PFR2, tapering off shortly before the flagellar tip (Figure 4C, 4D). Finally, we asked whether the PKAR1 localisation in this PFR domain depended on PFR2, PFC21 or PFR assembly factor 1 (Alves et al. 2020), which are required for formation of the outer PFR domain. Deletion of these genes in cells expressing PKAR1::mNG had no effect on PKAR1 localisation to the flagellum (Supplementary Figure Fig 5E-J). Together, these data suggest that flagellar PKAR1 is localised to the middle/inner PFR subdomain or at the PFR-axoneme connectors (PACs) with close proximity to doublet microtubules 4-7 (Sivadas et al. 2012; Zhang et al. 2022) (Figure 4E).

**Figure 4.**
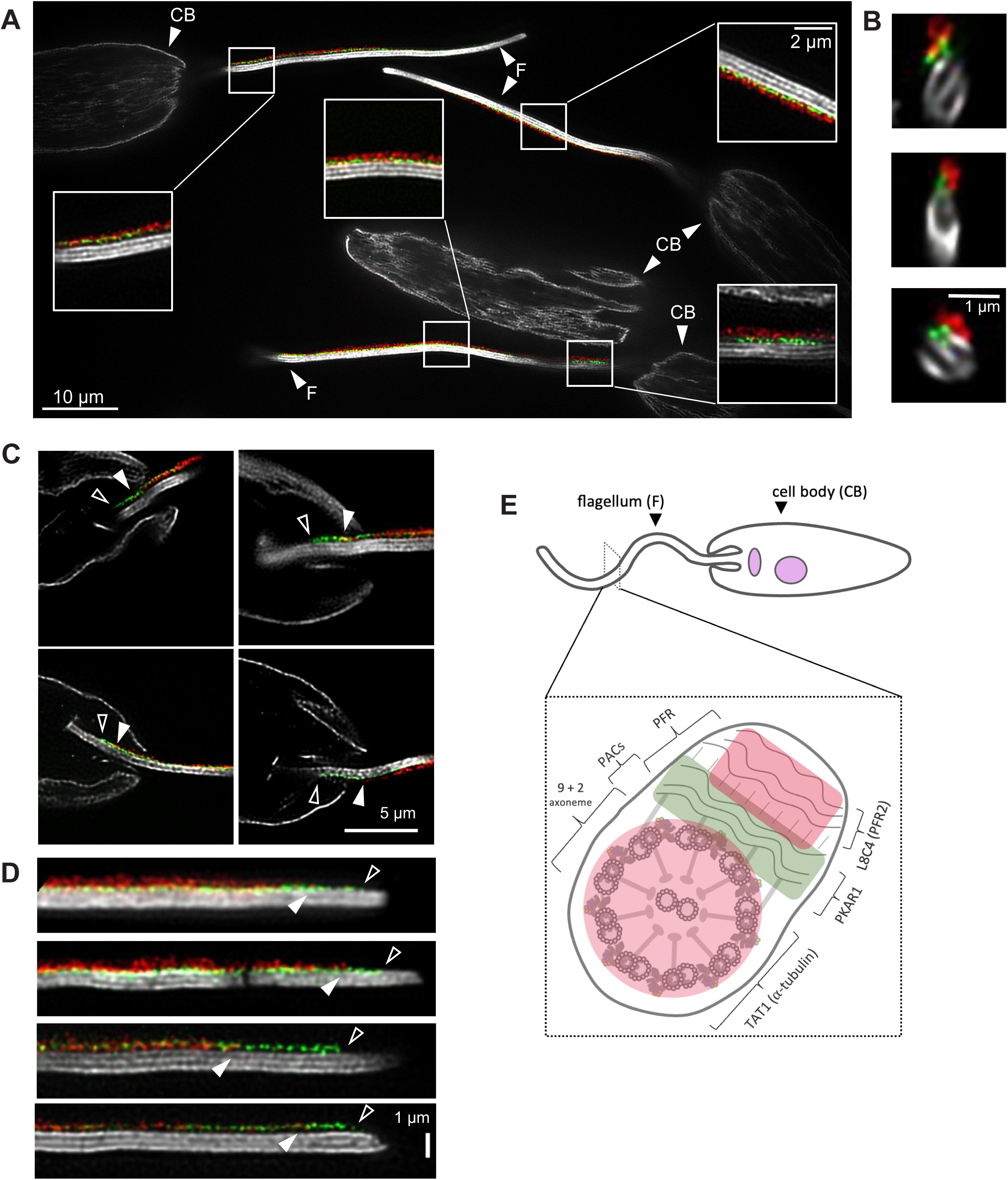
Localisation of LmxPKAR1 to the paraflagellar rod. Ultrastructure expansion microscopy of promastigotes expressing LmxPKAR1::mNG::c-myc. **(A)** A field of view depicting expanded promastigote cells with fluorescent signal corresponding to *Lmx*PKAR1 (anti-c-myc antibody, pseudocoloured green), PFR2 (anti-PFR2 antibody L8C4, pseudocoloured red), alpha-tubulin (anti-alpha-tubulin antibody, pseudocoloured grey). Insets show close-up views of flagellar segments where the LmxPKAR1 signal is seen between the PFR2 and tubulin signals. Abbreviations: CB, cell body; F, flagellum. **(B)** Flagellar cross sections showing the *Lmx*PKAR1 signal sandwiched between the PFR2 and tubulin signals. **(C)** Selection of micrographs showing the start of the *Lmx*PKAR1 signal before the PFR2 signal in the flagellar pocket. **(D)** Views showing the *Lmx*PKAR1 signal extending beyond the PFR2 signal in the distal flagellum. These were computationally straightened in ImageJ. Open arrows point to the distal end of the *Lmx*PKAR1 signal, filled arrows to the end of the PFR2 signal. **(E)** Cartoon of a promastigote flagellum cross-section showing the location of *Lmx*PKAR1, which may be anchored in the inner/middle domain of the PFR and/or at the PFR-axonemal connectors (PACs).

### Inosine dissociates PKAC1 signal from the flagellar cytoskeleton

To determine whether *Lmx*PKAR1 could be pharmacologically activated by purine nucleosides as was shown in *Trypanosoma* spp. (Bachmaier et al. 2019a; Ober et al. 2024), we used an indirect assay that measured solubilisation of mNG::PKAC1 from flagella treated with different compounds. Cytoskeletons from mNG::PKAC1 expressing cells were incubated with cAMP, toyocamycin, adenosine, guanosine and inosine prior to imaging and quantification of axonemal fluorescence (Figure 5).

**Figure 5.**
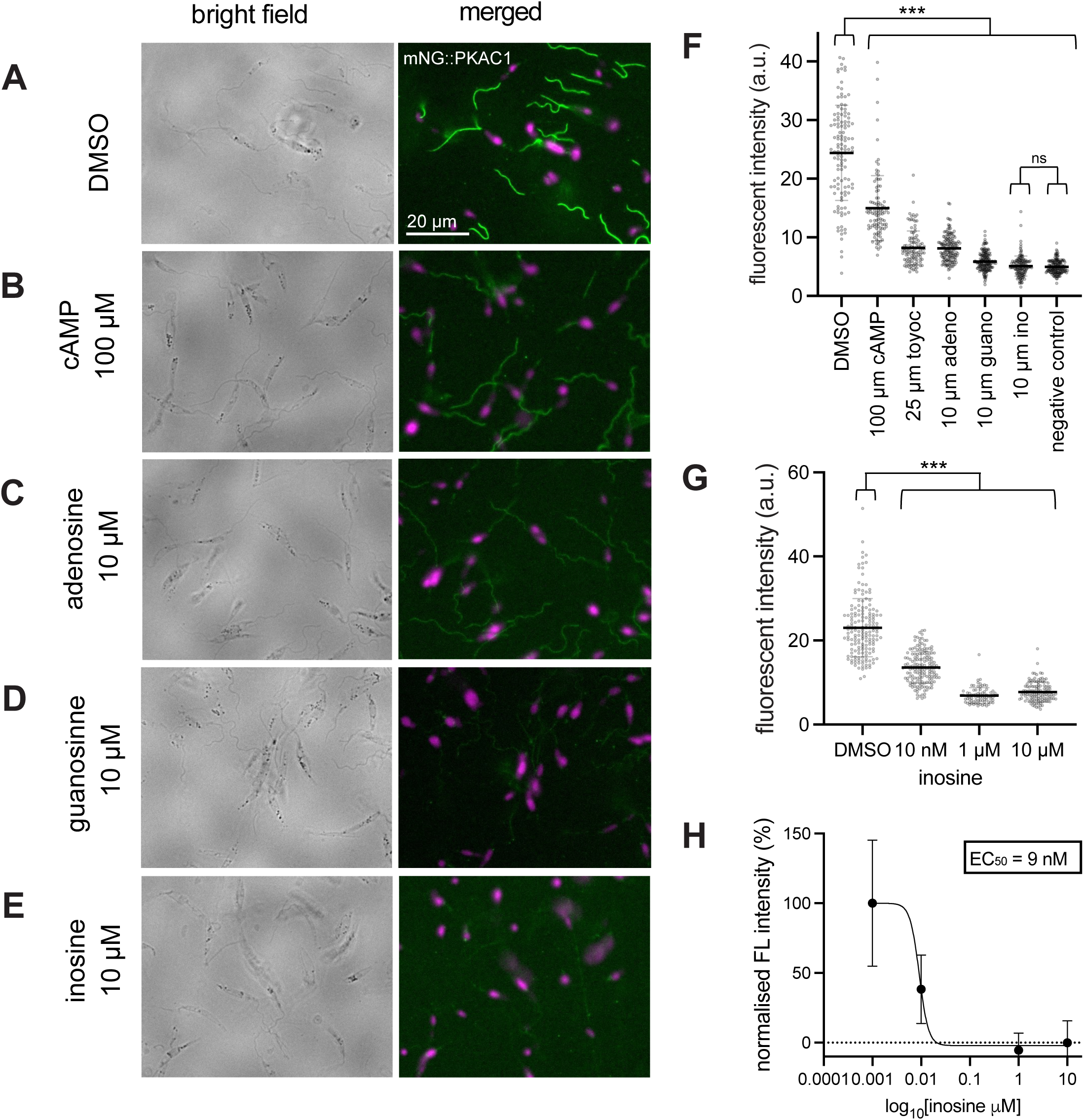
Pharmacological displacement of *Lmx*PKAC1 signal from the flagellum. Micrographs showing detergent-extracted promastigote cytoskeletons from mNG::*Lmx*PKAC1-expressing cells **(A)** without treatment (DMSO only control), **(B)** after treatment with 100 µM cAMP, and 10 µM **(C)** adenosine, **(D)** guanine and **(E)** inosine, respectively. Left panels, bright field images; right panels, merged fluorescence images of the mNG signal (green) and the signal from the DNA stain Hoechst (magenta). Scale bar, 20 µm. **(F)** Quantification of the cytoskeleton-associated mNG::*Lmx*PKAC1 signals from cells treated with 0.1% DMSO, 100 µM cAMP, 25 µM toyocamycin, 10 µM adenosine, 10 µM guanine, 10 µM inosine; untagged parental cells served as negative control. **(G)** Quantification of the cytoskeleton-associated mNG::*Lmx*PKAC1 signals in cells treated with inosine concentrations ranging from 10 µM to 10 nM. For all flagellar signal quantification, background fluorescence was subtracted from maximum intensity values. Each data point represents one cell measured across the flagellum midpoint. Horizontal bar indicates mean corrected fluorescent intensity. Significance was determined using an unpaired t-test assuming equal variance. *** represents a p-value < 0.001. n > 50 cell measured for each condition. **(H)** Normalisation of datapoints in (G) with a non-linear regression and calculated EC_50_. See Supplementary Table 4 for raw data.

The flagellar cytoskeleton mNG::PKAC1 signal remained visible after treatment with cAMP, at 100 µM (Figure 5A-B). However, quantification revealed a significant decrease in fluorescent signal relative to untreated mNG::PKAC1 cytoskeleton controls (Figure 4F). Toyocamycin was more efficacious at dissociating the mNG::PKAC1 signal at 25 µM than 100 µM cAMP (Figure 5F). Adenosine treatment at 10 µm had a similar effect on mNG::PKAC1 signal solubilisation than toyocamycin treatment. Guanosine and inosine were the most efficacious at reducing mNG::PKAC1 signal, with inosine treatment resulting in flagellar cytoskeletons with similar signal intensity to untagged negative controls (Figure 5C-F). Assessing a range of inosine concentrations on mNG::PKAC1 signal dissociation indicated an EC_50_ of ∼9 nM (Figure 5G-5H). This is consistent with recently published biochemical assays highlighting inosine as the highest affinity activator for trypanosomatid PKAR1, with guanosine and adenosine following at lower efficacies (Ober et al. 2024). Together, this adds to growing evidence that the physiological activating ligand is biochemically similar to purine nucleoside analogues like inosine, or perhaps inosine itself.

### LmxPKAC1 plays an important role in flagellar motility

We previously showed that partial deletion of *Lmx*PKAC1 (*ΔLmxM.34.4010*) reduced promastigote swimming speed (Beneke et al. 2019). Here, we employed the same CRISPR-Cas9 gene deletion strategy to generate the cell lines *ΔLmxPKAC1*, *ΔLmxPKAC2* and *ΔLmxPKAC3* as well as *ΔLmxPKAR1* and *ΔLmxPKAR3* (Supplementary Figure 6) to measure precisely how the loss of each of these proteins affected flagellar motility. The deletion mutants were all viable as promastigotes, with comparable *in vitro* growth rates and overall cell morphology as the parental cell line (Supplementary Figure 7). The swimming behaviour of deletion mutant populations was measured from darkfield videos (Wheeler 2017) and the results (Figure 6A, Supplementary Table 1) confirmed that promastigotes lacking *Lmx*PKAC1 swam significantly slower (average 4.9 μm/s ± 0.035 standard deviation, unpaired t-test assuming equal variance (p < 3.5 × 10^-14^) than the parental cell line (7.4 μm/s ± 0.917). *ΔLmxPKAC3* and *ΔLmxPKAR1* also swam slower (5.7 μm/s ± 0.032 and 5.4 μm/s ± 0.273, respectively); deletion of *LmxPKAR3* had the opposite effect and swam faster (8.4 μm/s ± 0.775). By contrast, deletion of axonemal *Lmx*PKAC2 did not significantly alter parasite swimming speed (average 6.4μm/s ± 0.650, unpaired t-test, p = 0.093) (Figure 6A). To test what the combined effect was of deleting *Lmx*PKAC1 and either *Lmx*PKAR3 or *Lmx*PKAC2 in the same cells, the respective double deletion mutants were generated (Supplementary Figure 6B). These showed the same reduced swimming speed as the *ΔLmxPKAC1* population (Figure 6A), indicating that loss of *Lmx*PKAC1 was the dominant cause of the observed slow motility phenotype, which could not be rescued or further compounded by secondary gene deletion of *Lmx*PKAR3 or *Lmx*PKAC2.

**Figure 6.**
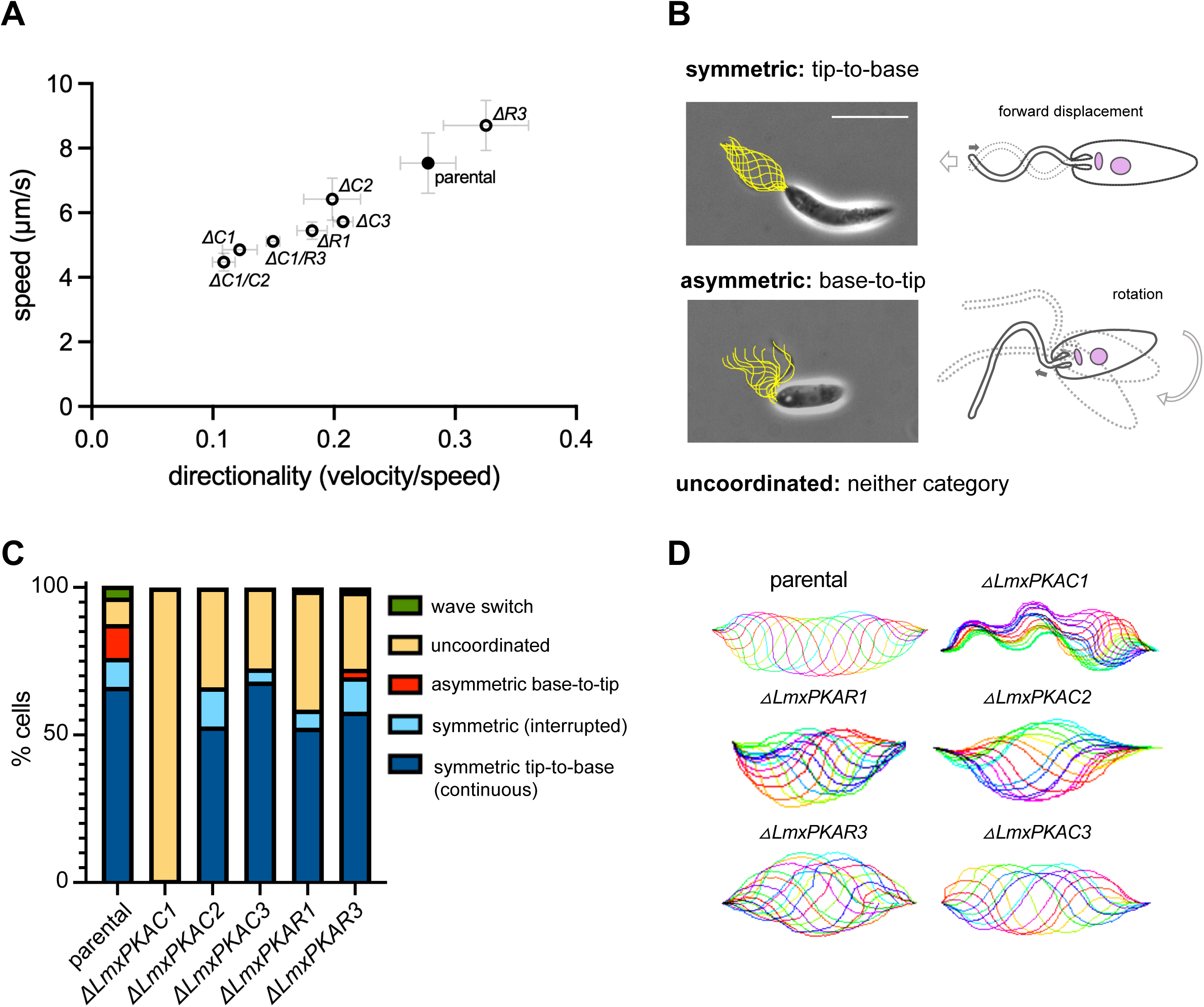
Mutant population motility. **(A)** Scatter plot of mean population swim speed and directionality (velocity/speed) of parental controls and deletion mutants for *Lmx*PKAC and *Lmx*PKAR subunits and of dual knock outs of *Lmx*PKAC1/PKAC2 and *Lmx*PKAC1/PKAR3. Swim speed measurement data from at least three replicates per population. Error bars indicate one standard deviation of the speed and directionality measurements. Raw data with p-values from unpaired t-tests for significance are summarised in Supplementary Table 1. **(B)** Phase contrast micrographs overlaid with real traces of flagellum position over one beat cycle, from a cell propagating either the (upper) symmetric tip-to-base flagellar wave for forward motility, or (lower) the asymmetric base-to-tip ciliary waveform for tumbling/turning. On the right are cartoon summaries of symmetric and asymmetric waves with a hollow arrow indicating the direction of cell displacement, and a filled grey arrow indicating direction of wave travel. **(C)** Relative instances of the waveforms observed in *ΔLmxPKAC* and *ΔLmxPKAR* mutants and parental control populations. **(D)** Computer-generated waveform reconstruction exemplified for each mutant, highlighting the lack of symmetry in △*LmxPKAC1* flagellar movement.

### LmxPKAC1 is required for the generation of symmetric flagellar waves

Since swimming speed measurements report the average for a whole population, we next used high speed video microscopy, alongside semi-automated tracing scripts (Walker, Ishimoto, and Wheeler 2019; Walker et al. 2019), to further dissect how deficiency of each PKA subunit was affecting the flagellar waveform of individual cells. First, we captured 0.5 seconds of high speed (200 Hz) low magnification (×20) videos to assess the relative proportions of the different waveforms that *L. mexicana* are known to generate: the tip-to-base symmetric flagellar waveform, the base-to-tip asymmetric ciliary waveform (Figure 6B), or, an aperiodic movement referred to as uncoordinated.

*ΔLmxPKAC3* and *ΔLmxPKAR3* showed similar proportions of cells performing symmetric tip-to-base waveforms as the parental cells (∼75%). However, all PKA deletion mutants appeared to display fewer instances of ciliary waveforms and a greater proportion of cells (> 25%) moving in an uncoordinated manner (Figure 6C). *ΔLmxPKAC2* and *ΔLmxPKAR1* both displayed a lesser proportion of cells propagating the flagellar wave (< 65%). Strikingly, *ΔLmxPKAC1* cells were completely unable to propagate the typical tip-to-base symmetric flagellar waveform and, instead, showed completely uncoordinated flagellum movements (Figure 6C, D, Supplementary movies 1 and 2). The uncoordinated movement and slow population swimming speed were rescuable by the addition of an episomal addback copy of *Lmx*PKAC1 (Supplementary Figure S8). This observation demonstrates that *Lmx*PKAC1 is essential for producing a symmetric and rhythmic flagellar waveform.

### All PKAC subunits contribute in different ways to beat amplitude and frequency

In order to dissect how the loss or dysregulation of PKA contributes to the overall waveform, we used cell tracing and Fourier analysis to extract the dominant beat frequency, amplitude and waves per flagellum comprising the tip-to-base symmetric waves for the parental control cells each PKA deletion cell line (Figure 7A-F). *L. mexicana* tip-to-base waves usually have a frequency of 20-35 Hz and an amplitude between 40 to 60 degrees (Walker et al. 2019; Wang et al. 2020). This was confirmed here for the parental control cell line (Figure 7, p-values listed in Supplementary Table 2). *ΔLmxPKAC1* were never observed propagating a rhythmic symmetric flagellar wave (Figure 6C). Our best effort to fit a dominant frequency of attempted tip-to-base movements showed a greatly diminished frequency (< 10 Hz), with an average shear angle amplitude of 26 degrees (Figure 7A), indicating that *Lmx*PKAC1 is required for the generation of a normal flagellar waveform. The flagellar length and wavelength are coupled in *Leishmania* and relates species, leading to tendency of ∼1 wavelength per flagellum (Gadelha, Wickstead, and Gull 2007). We saw here that these aforementioned parameters showed a positive correlation in all PKA deletion cell lines, except *ΔLmxPKAC1* cells (Supplementary Figure 9). Flagellar waves from *ΔLmxPKAC2* displayed a slight but significantly diminished beat amplitude (Figure 7B). We also observed that *ΔLmxPKAC3* had significantly diminished beat frequencies (averaging 22 Hz), but otherwise displayed normal beat amplitudes (Figure 7C). *ΔLmxPKAR1* retained comparable flagellar wave parameters relative to parental control cells (Figure 7D). Lastly, although *ΔLmxPKAR3* and *ΔLmxPKAC2* mutants showed a comparable average beat frequency for tip-to-base symmetric waves to parental control populations, the measured values appear to show a bimodal distribution, with a sub-population with low beat frequency (∼15 Hz) and one with an abnormally high beat frequency (> 40Hz) (Figure 7B, 7E). The normal average population speed for *ΔLmxPKAC2* and elevated population swimming speed for *ΔLmxPKAR3* (Figure 6A) could have arisen from the averaging of measurements from a population displaying these extreme frequency values. More cells would need to be captured, and perhaps observed over longer time periods, to ascertain the true relative proportions of these phenotypes in the population and assess whether individual flagella can switch between abnormally low and high frequencies.

**Figure 7.**
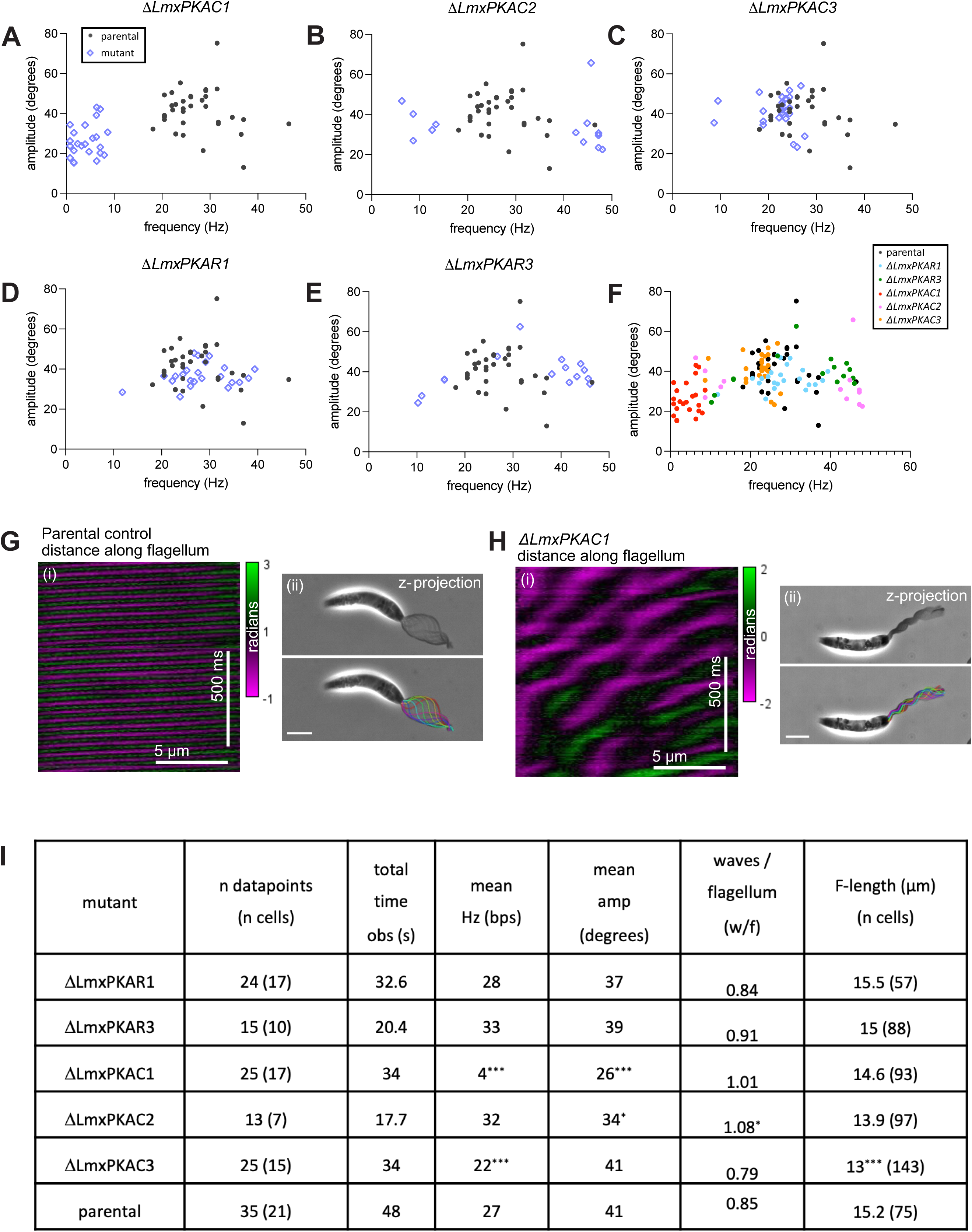
Symmetric wave amplitude and frequency parameters. Fourier decomposition of the tip-to-base uncoordinated wave attempts of △*LmxPKAC1,* and the symmetric flagellar waves of △*LmxPKAC2,* △*LmxPKAC3,* △*LmxPKAR1 and* △*LmxPKAR3.* **(A-E)** Scatter plots of the dominant beat frequency in hertz (Hz) plotted against beat amplitude in degrees for **(A)** △*LmxPKAC1,* **(B)** △*LmxPKAC2,* **(C)** △*LmxPKAC3,* **(D)** △*LmxPKAR1* **(E)** △*LmxPKAR3* populations (purple diamonds) overlaid with measurements from parental control cells (black filled circles). Each point represents one unit of time observed (see Methods). **(F)** All datapoints across (A-E) representing beat frequency and amplitude represented in one scatter plot. **(G)** Tangent kymograph (i) and flagellar traces (ii) of a parental control cell; distance is on the x-axis, time one the y-axis. **(H)** Tangent kymograph (i) and flagellar traces (ii) of a *ΔLmxPKAC1* cell. Tangent kymographs follow 1 second of flagellar beating, and z-projections of 150 ms of beating with overlayed traces of flagellum position every 25 ms where rainbow coloration indicates start (red) to end (indigo) of projected time. **(I)** Summary numbers for each deletion mutant cell line: Number (n) of timepoints captured by high-speed video microscopy, where each timepoint represents 256 video frames, and n cells captured in total; the total time observed for each mutant dataset in seconds. The mean angular amplitude (amp) of the waveform in degrees (deg); mean beat frequency (beats per second, bps; Hz); number of waves per flagellum (w/f); flagellum length in µm and number (n) of cells for which length was measured. Mann-Whitney **U** test was used to ascertain significance of each dataset against that of the parental control measurements. Significant p-values are indicated as, *** = p < 0.001, ** = p < 0.005, * = p < 0.05. All p-values can be found in Supplementary Figure 7. See Supplementary Table 5 for raw data.

Taken together, these measurements show that removal of different PKA subunits perturbed the flagellar waveforms of *L. mexicana* in specific ways. Removal of the catalytic subunit PKAC1, which is normally docked to the flagellar cytoskeleton through association with PKAR1, had the biggest effect; without PKAC1 the flagellar waveforms lost their symmetry and this uncoordinated beating reduced the speed at which the cells swam.

## Discussion

Protein kinase A is one of the most studied eukaryotic signal transducers, which is highly conserved throughout the eukaryotes (with the exception of most plant lineages) (Peng et al. 2015a) and fine-tunes a multitude of cellular processes. Within motile cilia, PKA activity modulates sperm beating behaviours, capacitation and hyperactivation, which are both essential for reproduction (Alonso et al. 2017; Lefièvre et al. 2002). Despite this importance, there is a lack of clear understanding of how PKA activity is spatially controlled and how its activity fine-tunes flagellar waveform properties. In kinetoplastids, converging evidence from proteomics and transcriptomics datasets and mutant phenotypes suggested their divergent PKA may also have a role in flagellar motility regulation. Here, we used a genetic approach combined with fluorescence-, expansion- and video-microscopy, to dissect the localisations and paring preferences of the three *L. mexicana* PKAC and two PKAR subunits and studied how gene deletions impacted cell swimming and generation of symmetric waveforms. Our results corroborate recent reports of PKAC/R pairings in *Leishmania* (Fischer Weinberger et al. 2024) and extend previous analyses on PKA function by providing multiple lines of evidence for a necessary function for *Lmx*PKAC1 in flagellar waveform regulation.

We found that there are two regions of PKA concentration in promastigote forms: *Lmx*PKAC1, *Lmx*PKAC2 and *Lmx*PKAR1 showed enrichment in the flagellum, while a strong cell body cortex signal was detected for *Lmx*PKAC3 and *Lmx*PKAR3. These localisations were neither exclusive nor static and examination of cells at different stages in the division cycle indicated relocalisation of all three *Lmx*PKAC subunits and of *Lmx*PKAR3; only *Lmx*PKAR1 maintained its dominant flagellar signal in all examined cell cycle stages, and withstood detergent extraction. Upon deletion of *Lmx*PKAR1, the flagellar signal of *Lmx*PKAC1 was greatly diminished and we conclude that *Lmx*PKAR1-*Lmx*PKAC1 interaction is necessary for *Lmx*PKAC1 localisation to the flagellum.

Characterising the phenotypes of gene deletion mutants showed that *Lmx*PKAC1 was essential for propagation of symmetric flagellar waveforms used for forward motility. All PKA deletion cell lines were viable and remained motile (including the double deletion of *Lmx*PKAC1 and *Lmx*PKAC2), but measurement of swimming speeds showed differences compared to the parental control line for all deletion mutants except *ΔLmxPKAC2* (Figure 6A). *ΔLmxPKAC1* showed the greatest reduction in swimming speed and this mutant was unique in its complete inability to generate any symmetric tip-to-base flagellar waves (Figure 6C), instead beating its flagella in an uncoordinated manner characterised by a lack of periodicity, symmetry, a very low dominant beat frequency and reduced amplitude compared to normal tip-to-base waveforms of promastigotes (Figure 7). These data suggest *Lmx*PKAC1 is required within the flagellum to maintain the appropriate spatio-temporal activation of axonemal dyneins, most likely as a component of a signal transduction cascade that leads to the phosphorylation of either core axonemal proteins that regulate dyneins, or of dynein subunits directly. The ODAs, which are important for the frequency of base-to-tip asymmetric waveform in *Chlamydomonas* (Brokaw and Kamiya 1987), are thought to be a key target of cAMP-dependent protein kinase (PKA) (models, reviewed in (Tash 1989; Cheng, Zhang, and McCammon 2006)). Outer dynein light chain proteins from the ciliates *Paramecium tetraurelia* (p29 (Barkalow, Hamasaki, and Satir 1994)) and *Tetrahymena* (p34 (Christensen et al. 2001)) were shown to be phosphorylated upon treatment with cAMP and RNAi knockdown of *P. tetraurelia* outer dynein light chain 1 (ODLC1) resulted in loss of phosphorylated p29 and rendered cilia unresponsive to cAMP (Kutomi et al. 2012). It will be interesting to test whether ODAs are targets for the divergent cAMP-independent PKA in kinetoplastids. Phosphoproteomic data from *T. brucei* treated with a synthetic PKA activator, 7-CN-7-C-ino, identified phosphopeptides for several proteins with known functions in normal motility including several dynein intermediate chains (e.g. IC138), distal docking complex 2 (dDC2) and nexin dynein regulatory complex-2 (DRC2) (Bachmaier et al. 2019a). The specificity and functional significance of any of these phosphosites remains to be tested. Comparisons of phosphorylation patterns in axonemes of *L. mexicana* PKA mutants and wild type controls could further corroborate candidates for *Lmx*PKAC1 phosphorylation substrates and allow for further dissection of the mechanism by which phosphorylation contributes to flagellar waveform modulation.

Cells lacking *Lmx*PKAR1 also swam significantly slower, and this mutant lost most of its flagellar *Lmx*PKAC1 signal. This is consistent with a model where axonemal *Lmx*PKAR1 binds and concentrates PKAC1 along the flagellum, placing it in proximity to axonemal proteins which are phosphorylated upon receipt of an activating signal. The expansion microscopy data place the *Lmx*PKAR1 at the interface between axoneme and PFR structure (Figure 4). This suggests a model where *Lmx*PKAC1 activity is concentrated asymmetrically within the motile axoneme with proximity to doublets 4-7. It is tempting to speculate that the different placement of PKAR in different types of motile cilia is related to their distinct waveforms and preferred wave propagation direction: localisation to distinct radial spokes is observed in *Chlamydomonas* and mammalian cilia (Leung et al. 2025), with exclusively base-to-tip propagation, while the asymmetric extra-axonemal PKAR localisation in kinetoplastids occurs in flagella with a need to modulate both near-planar tip-to-base and asymmetric base-to-tip beat types (Gadelha, Wickstead, and Gull 2007; Wang et al. 2020). *Leishmania* lacking the outer PFR domain due to deletion of PFR2 swim slower (Santrich et al. 1997) but still propagate predominantly tip-to-base waveforms that tend to remain near symmetric, with variable frequency and lower amplitude (Wang et al. 2020). These mutants do not fully phenocopy the defects in the *ΔLmxPKAC1* mutants. Deletion of PFR assembly factors or major components of the outer and middle/inner domains (PFR2 array and PFC21, respectively) leave a remnant PFR structure intact (Santrich et al. 1997; Alves et al. 2020; Gabriel et al. 2024), which still retains PKA in *Leishmania* (Supplementary Figure 5E-I). Future identification of the PAC components necessary for tethering the PFR to the axoneme could provide a route to a clean removal of the entire PFR structure, which would allow further studies to observe the *Leishmania* wave shape upon complete removal of the asymmetric PFR structure.

Of course, it is not unusual for kinases to have pleotropic function and moreover, localisation of proteins does not necessarily indicate function within that domain. Likewise with the observed dual localisation of *Lmx*PKAC1, its localisation in the flagellum in non-dividing cells could be, in part, a sequestration away from other targets within the cell body or nucleus, which may be involved in replication events. However, while the mutant analysis revealed a clear motility phenotype upon loss of *Lmx*PKAC1, cell growth remained unaffected, arguing against a key role in the regulation of cell division. Alternatively, during division, where forward swimming may need to be dampened, *Lmx*PKAC1 may be partitioned to the nucleus until returning to the new and old flagella. Experiments involving pulse-chase labelling of proteins would be required to assess the relationship between the different *Lmx*PKAC1 populations fractions over the course of the cell cycle. In any case, the activation status of *Lmx*PKAC1 in the nucleus remains an open question. *Lmx*PKAR1 was never observed alongside its catalytic subunit in the nucleolus during division and this spatial uncoupling would presumably populate the nucleolus with active *Lmx*PKAC1 or, if sequestration was the purpose, require an alternative regulatory mechanism for *Lmx*PKAC1 inactivation.

*Lmx*PKAC1 and *Lmx*PKAC2 share 86% sequence identity and localisation data suggest *Lmx*PKAR1 is required for their enrichment in the flagellum. Unlike *Lmx*PKAC1, however, *Lmx*PKAC2 was solubilised by treatment with 0.1% NP-40 while *Lmx*PKAC1 remained in the flagellar cytoskeletons. Furthermore, loss of *Lmx*PKAC2 had only a small effect on flagellar waveforms when observed at the level of populations: *ΔLmxPKAC2* displayed comparable population swimming speeds to parental controls, with only a small increase in the proportion of cells exhibiting uncoordinated flagellar waveforms (Figure 6C). The observation that the frequencies of symmetric waveforms of individual cells segregated into two groups, low frequency or high frequency, suggests that *Lmx*PKAC2 may have a role in maintaining a constant beat frequency (Figure 7B). The population overview waveform analyses represent snapshot observations of flagella over a short time period (0.5 seconds). To study this further, it would be informative to observe individual cells for a longer time to determine if periods of low frequency waves alternate with high frequency waves in single flagella.

*Lmx*PKAC3 had a clearly distinct localisation, concentrated at the cell cortex, similar to *Lmx*PKAR3. The peripheral enrichment was lost upon deletion of *Lmx*PKAR3, suggestive of a *Lmx*PKAC3/R3 complex. A faint flagellar signal was also observed for *Lmx*PKAC3, which was increased in the *ΔLmxPKAR3* deletion mutant. Interestingly, the motility phenotype of gene deletion mutants correlated with changes of *Lmx*PKAC3 abundance in the flagellum: the *ΔLmxPKAC3* population swam more slowly and *ΔLmxPKAR3* swam faster than parental controls. The slower swimming speed for *ΔLmxPKAC3* was likely the combined effect of the increased proportion of cells showing uncoordinated waveforms and the significant decrease in the beat frequency of symmetric waves (Figure 7C). Additionally, there was a small reduction in average flagellum length (Supplementary Figure 7D). Interestingly, loss of *Lmx*PKAR3, which led to a re-distribution of *Lmx*PKAC3 away from the cell cortex, resulted in more cells propagating flagellar waveforms exceeding 40 Hz (Figure 7E) and increased mean population swimming speed (Figure 6A). It remains an open question whether *Lmx*PKAC3 has bona-fide phosphorylation targets within the flagellum, or whether the increased amount of soluble catalytic subunit in the *Lmx*PKAR3 deletion mutant causes off-target phosphorylation events in the axoneme that affect motility, secondary to its main role elsewhere. As deletion of *Lmx*PKAR3 (shown to increase flagellar *Lmx*PKAC3 Figure 3B) in *ΔLmxPKAC1* flagella did not rescue the motility defect of *ΔLmxPKAC1* (Figure 5A), it seems probable that these catalytic subunits have distinct phosphorylation substrates within the motile flagellum. A recent report linked *Lmx*PKAR3 and *Lmx*PKAC3 to processes involved in cell morphology in *L. donovani* (Fischer Weinberger et al. 2024). This may explain the shortened flagella in our *ΔLmxPKAC3* cells; however, we did not observe any morphological changes in *ΔLmxPKAR3*. Moreover, all of our *L. mexicana* PKA deletion mutant cell lines here showed normal cell growth and maintained the same mean cell pseudo-diameter as the parental controls (Supplementary Figure 7).

Owing to the high conservation of PKA, it has long been presumed that its activation mechanism through cAMP was equally as conserved, until the recent discovery that trypanosome PKA is insensitive to cAMP and instead activated by purine nucleoside analogues (Ober et al. 2024; Bachmaier et al. 2019a). Multiple lines of evidence indicate that this alternative PKA activation pathway also operates in *L. mexicana*: The sequences of the *Lmx*PKAR1 and *Lmx*PKAR3 phosphate binding cassettes showed some of the characteristic residue switches that were shown in *Trypanosoma* to prevent cAMP binding. Moreover, treatment of cytoskeletons with cAMP or nucleosides to test for dissociation of the *Lmx*PKAC1 signal revealed inosine as the most potent (EG_50_ 9 nM), followed closely by guanosine (Figure 5). This is an indirect assay for PKA activation and the results are consistent with the model of the *T. brucei* PKAR1/C2 complex, where inosine binding to PKAR resulted in the release of the catalytic subunit and subsequent substrate phosphorylation (Bachmaier et al. 2019a; Ober et al. 2024). As inosine is an intermediate compound of the purine metabolism pathway (Boitz et al. 2012), it may be that the endogenous ligand for divergent PKA signalling is a derivative of this pathway, or inosine itself.

Although the role of cAMP-responsive PKA in flagellar signalling has been established (Soriano-Úbeda et al. 2019; Lefièvre et al. 2002), detailed structural maps of ciliary axonemes revealed variations in PKA positioning between *Chlamydomonas* and humans. Our study adds to this emerging picture of lineage-specific positional variation, and shows that second messengers outside of cAMP and calcium may be used for flagellar waveform modulation through PKA. Despite this diversity of signal transducers, it will be interesting to discover whether the signals converge at conserved regulatory structures to orchestrate the spatio-temporal activity of axonemal dyneins.

## Methods

### Cell culture

All cell lines used in this project were derived from *Leishmania mexicana* (WHO strain MNYC/BZ/62/M379) cell line expressing Cas9 and T7 RNA polymerase (Beneke et al. 2017). Promastigote form cells were cultured exactly as described in (Beneke et al. 2017). In brief, cells were incubated at 28°C and maintained in M199 (Life Technologies) medium supplemented with 2.2 g/L sodium bicarbonate between 1×10^5^ and 2×10^7^ cells/ml with regular sub-culturing.

### Endogenous fluorescent tagging and gene deletion

Gene deletions and cell lines expressing endogenously fused fluorescent gene products were essentially created as outlined by Beneke *et al*., (Beneke et al. 2017). An online resource for automated primer design (www.LeishGEdit.net) was used to obtain primer sequences for the replacement of target gene ORFs with drug resistance cassettes (pT plasmids used to create donor products) or insertion of mNeonGreen (pPLOT plasmids used to create donor products) to the endogenous gene locus (Beneke et al. 2017). Promastigote populations transfected with DNA donor and guide constructs were maintained in M199 medium supplemented with relevant selection drugs (either 20 µg/ml puromycin, 5 µg/ml Blasticidin or 40 µg/ml G418) for 3 culture passages before performing a PCR based gene deletion verification.

### Diagnostic PCR for gene deletion validation

Primers sequences targeting the open reading frames (ORF) of genes of interest (GOI) were taken from the LeishGEdit primer database (www.LeishGEdit.net) and sourced from ThermoFisher. Due to high sequence similarity, *LmxPKAC1* and *LmxPKAC2* gene deletion verification, primers were manually designed using Primer3 (see Supplementary Table 3A for primer sequences and corresponding PCR amplicon sizes). To test for presence or absence of a GOI, the relevant forward and reverse (F/R) primers were used to amplify a fragment of the ORF from genomic DNA from the parental cell line (positive control) and genomic DNA from the deletion mutant. A PCR reaction targeting a “housekeeping” gene, here PF16 (LmxM.20.1400) or IFT88 (LmxM.27.1130), was run in parallel as a technical control. Master mixes for the PCR reactions constituted the FastGene Optima HotStart Ready Mix sourced from Nippon Genetics containing 4 ng/μl of the relevant gDNA and 2.5 μM each of the forward and reverse primers. For all diagnostic PCRs, the following PCR program steps were used. Step 1: First denaturation at 95°C for 3 minutes. Step 2: denaturation at 95°C for 15 seconds. Step 3: 58°C for 15 seconds. Step 4: 72°C for 30 seconds. Step 5: repeat steps 2-4 for 30 cycles. Step 6: 72°C for 1 minute.

### Detergent extraction of cell membranes

Promastigote cytoskeletons were prepared from 1 – 5 ml (depending on the cell culture density assessed by eye) of mid-log cells. Cells were washed once with PBS and the washed pellet was resuspended in 100 µl (or 1 ml if > 1E7 cells are in the sample) PEME [100 mM Pipes, 2 mM EGTA, 1 mM MgSO4 and 100 nM EDTA] with 0.1% NP-40. Once homogenous and the solution turns clear, the sample was spun at 2°C at 10,000*g* for 5 minutes. The supernatant was removed and the pellets were washed with PEME and then resuspended in PEME or PBS with 10 µg/ml Hoechst 33342. For the experiments assessing the fluorescent *Lmx*PKAC1 signal in cytoskeletons, the treatment substance was added to the PEME used during the final wash step at a 1:1000 ratio from a stock solution at 1000x final concentration (cAMP (Sigma-Aldrich A6885), at 1 and 100 mM in water; toyocamycin (Sigma-Aldrich T3580) at 1 and 100 mM in DMSO; inosine (ThermoFisher Scientific A14459); 1 and 100 mM in water). If the substance was dissolved in DMSO, then the respective amount of DMSO (0.1%) was added to the control cytoskeletons in PEME buffer.

### Light microscopy

For live cell fluorescence microscopy of promastigote cells, 1 ml of a mid-logarithmic culture (between 1×10^6^ – 1×10^7^ cells/ml) was spun at 800*g* for 5 minutes and the pellet resuspended in 100 µl of PBS and [10 µg/ml] Hoechst 33342 (ThermoFischer). A volume of 0.7 µl was spread onto a poly-lysine coated glass slide, bordered (∼ 5 x 2 cm rectangle) with a hydrophobic fat pen (Dako). A glass coverslip, thickness 1 or 1.5 was used to cover the slide. Live cell imaging was typically done with a Zeiss Axioimager.Z2 microscope with a Hamamatsu ORCA-Flash4.0 camera, x63 oil objective lens with numerical aperture (NA) 1.4, and phase contrast illumination. Fluorescence channel was acquired with between 0.5 and 3 seconds of excitation depending on the brightness of the fluorescent signal.

For live cell imaging of axenic amastigotes, 500 µl of mid-log promastigote culture was inoculated into a vented 25 cm^3^ flask (total 5 ml volume) containing Schneider’s Drosophila medium (Life Technologies) supplemented with 20% heat-inactivated foetal bovine serum (FBS) (Gibco) and adjusted to pH 5.5 with concentrated HCl. Cultures were incubated at 34°C with 5% CO_2_ for 72 hours prior to imaging. 1 ml of sample culture was pelleted by centrifugation at 800*g* for 5 minutes, pellet washed with PBS, spun again, then the washed pellet resuspended in 50 µl of PBS plus Hoechst 33342 (final 10 µg/ml). A volume of 20 µl was placed on the poly-lysine coated glass slide and left inside a humid chamber for 10 minutes to settle. Excess liquid was gently removed by pipetting and the cover slide placed on top.

### Expansion microscopy

The protocol for ultrastructural expansion microscopy was adapted from (Gambarotto, Hamel, and Guichard 2021). Five million exponentially growing *Leishmania* promastigotes were washed twice in PBS before settling on a 12 millimetre glass coverslip in a 24-well plate for 10 minutes. Excess liquid was removed, and cells were fixed with 2 % formaldehyde (FA) (ThermoFisher, 28906) for 10 minutes. Samples were permeabilized using 0.1% Triton X-100 for 1 minute before washing twice with PBS. Samples were blocked with 2% blocking buffer (2% bovine albumin in PBS) for 10 minutes with gentle shaking before being incubated with primary antibodies (Supplementary Table 3B) in blocking buffer overnight at room temperature. After three washes with 0.1% Tween in PBS (PBS-T), samples were incubated with secondary antibodies in blocking buffer for 2 hours at room temperature with gentle shaking, and all subsequent steps were performed in the dark. For samples destined for microscopy quality control check, Hoechst 33342 (Thermofisher 62249) was added with the secondary antibody at 20 µg/ml. After staining was confirmed to be high quality and specific (i.e. no corresponding signal observed in the untagged parental control), samples were anchored in 2% acrylamide (Merck, A4058), 1.4 % FA for 2 hours at 37 °C before being washed with PBS. Sample gelation was performed in an ice-filled humid chamber on top of a parafilm layer. Ammonium persulfate (APS) (Merk, A3678) and N,N,Ń,Ń-tetramethylethylenediamine (TEMED) (Merk, T9281) were added to monomer solution to a final concentration of 0.5 % each, vortexed briefly, then the sample coverslip placed sample-side-down onto 35 µl gelation solution atop the parafilm. After 5 minutes incubation on ice, samples were transferred to 37 °C for 30 minutes before removed from the parafilm and added to 1 ml of denaturation buffer (200 mM SDS, 200 mM NaCl, 50 mM Tris in water, balanced to pH = 9 using NaOH) in a 6 well-plate. After 10 minutes of gentle shaking, the glass coverslips were discarded and the gelated sample transferred to a 2 ml Eppendorf with 1 ml fresh denaturation buffer and heated to 95 °C for 30 minutes. The gel was washed thrice in 5 ml PBS before a quarter was cut out and incubated at room temperature overnight in guineapig anti alpha-tubulin primary antibody (ABCD Antibodies, ABCD_AA345) in blocking buffer for post staining. The gel sample was then washed thrice in PBS-T then incubated for 2 hours at room temperature with the corresponding anti guineapig secondary antibody (Fisher Scientific, 10624773) in blocking buffer. A final three washes with PBS were done prior to placing the sample in 50 ml of Mili-Q water. The water was changed every 20 minutes until the gel had reached maximum expansion (estimated to be ∼4.5 fold based on axoneme width). Lastly the gel was mounted onto a poly-L-lysine coated round sample holder and imaged with a Nikon Ti2 Cicero spinning disk microscope using the 100X objective (NA 1.45) and the Kinetix sCMOS Camera. Images were further deconvolved using Huygens software version 3.10 (Scientific Volume Imaging, The Netherlands, http://svi.nl) using the following parameters (Acuity: 0, SNR:28,27,20, 30 iterations, background estimation: auto), then the maximum intensity z-projection taken unless otherwise indicated.

### Cell swimming speed and directionality

Cell swimming speed and directionality were measured from 512 frame-long darkfield videos of cell cultures in a 0.1 mm deep chamber, captured at 5 frames per second (fps) using a ZeissAxioimager.Z2 microscope with 10× objective and an Andor Neo 5.5 camera, exactly as described in (Wheeler 2017) with the modification described in (Beneke et al. 2019).

### Population overview waveform analysis (POWA)

To quantify the proportion of cells undergoing different waveforms, 1 ml of exponential growth phase promastigote cells (1×10^6^ – 1×10^7^ cells/ml) was centrifuged for 5 minutes at 800*g*. Between 600 – 900 µl supernatant was removed and the cells resuspended in the remaining 400 – 100 µl of M199 medium, effectively concentrating the cells. 1 µl of 1:100 bead mix (Sigma 79633 polystyrene microparticle beads (5 µm size) in M199 medium) was added to cell suspension and mixed gently until homogenous. 1 µl of cell sample was added to the centre of a glass microscope slide bordered by a Dako hydrophobic pen (∼5 cm by 2 cm rectangle). A glass coverslip of 1.0 or 1.5 thickness bordered by a similar Dako hydrophobic pen rectangle was placed gently on top of the sample slide, effectively creating a sealed chamber. Videomicrographs of swimming cells under phase contrast illumination were captured with an Andor Neo 5.5 camera at 200 frames/s for 0.5 seconds, using a ×20 NA 0.3 objective lens on a Leica Inverted microscope. Manual categorisation of waveforms was performed in ImageJ or Fiji (Collins 2007; Schindelin et al. 2012) for non-dividing cells only. Categories include symmetric wave tip-to-base (continuous and interrupted), asymmetric wave base-to-tip and uncoordinated which is neither symmetric tip-to-base nor asymmetric base-to-tip. Flagellar length was measured from stills obtained from these videomicrographs, using the Fiji freehand line measuring tool.

### Waveform parameter analysis

The capture of videomicrographs for waveform analysis was exactly as above (POWA), with the exception of using a ×100 NA 1.4 oil immersion objective and capturing for 5 seconds duration. All videomicrographs were captured within 10 minutes of making sample slides. Tip-to-base beat waveforms were analysed in ImageJ, using custom scripts (https://github.com/zephyris/flagellar-beat-fourier-ij). Flagella were traced using the midline method as described in (Walker, Ishimoto, and Wheeler 2019), except the cell body was manually identified. Data was represented as a tangent angle kymograph, plotting tangent angle at each position in the flagellum over time. The resulting tangent-angle kymographs were manually cropped to a length of either 128 (0.64 seconds) or 256 (1.28 seconds) frames. Each datapoint on the plots represent one kymograph (cropped to the lengths described above), where several datapoints (up to three) could derive from one cell.

Tangent angle kymographs were analysed using a Fourier transform-based method to extract beat parameters, using the concepts described in (Walker et al. 2019). The Fourier power spectrum of the tangent angle was calculated and summed over the length of the flagellum. The dominant beat frequency was obtained by determining the frequency with the greatest power from the power spectrum using a sliding window with a width of 10% of the measurable frequency range.

Data obtained from Fourier analysis were compiled in Excel and plotted using GraphPad Prism 9. Amplitude measurements along each flagellum (computed for the identified dominant frequency from the FFT and defined in terms of the tangent angle) were averaged and plotted against the dominant frequency. The number of complete sinusoidal waves present along the flagellum (equivalent to the wavelength divided by the flagellum length) was calculated in Excel by taking the difference in phase of the identified dominant frequency between the proximal and distal ends of the flagellum, then dividing the phase difference by 2π. This analysis effectively approximates the flagellar waveform as a single sinusoidal mode (when expressed in terms of tangent angle), as justified by observations of the wild type (Walker et al. 2019).

## Supporting information

Supplementary_movie1_parental_20X 100fps 1s_014-2_quarter_speed_playback

Supplementary_movie2_PKAC1KO_20X_140fps_1s-1_quarter_speed_playback.tif

Supplementary_Table1_population_motility_raw_data

Supplementary_Table2_statistical_test_waveform

Supplementary_Table3_primers_antibodies

Supplementary_Table4_rawdata_cytoskeleton_dissociation

Supplementary_Table5_waveform_parameters_raw_numbers

## Acknowledgements

A part of this work was carried out at the Wellcome Centre for Integrative Parasitology (WCIP) at the University of Glasgow and we thank the WCIP for hosting SF as a visiting student. We thank Keith Gull (University of Oxford) for the gift of antibody L8C4 and access to laboratory equipment at the Dunn School, Clirim Jetishi (Ochsenreiter lab, University of Bern) for help with ultrastructure expansion microscopy and Michael Boshart for stimulating discussions.

