## Supplementary figures and images for "Divergent Protein Kinase A contributes to the regulation of flagellar waveforms in *Leishmania mexicana*"

### Supplementary_movie1_parental_20X 100fps 1s_014-2_quarter_speed_playback

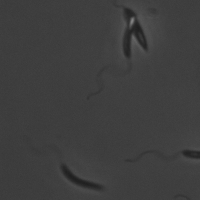

### Supplementary_movie2_PKAC1KO_20X_140fps_1s-1_quarter_speed_playback.tif

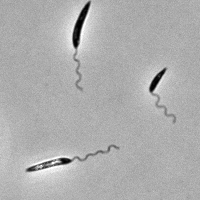
