## Supplementary_Table2_statistical_test_waveform for "Divergent Protein Kinase A contributes to the regulation of flagellar waveforms in *Leishmania mexicana*"

**Supplementary Table 2.** Waveform parameters from mutants and the parental control populations were compared with a Mann-Whitney U test. The table shows the p-values, with significance indicated by asterisks as follows, *** = p < 0.001, ** = p < 0.005, * = p < 0.05. Datasets analysed include wave amplitude, dominant frequency (Hz), waves per flagellum (w/f) and flagellum length (F-length).

| | ***ΔLmxPKAR1*** | ***ΔLmxPKAR3*** | ***ΔLmxPKAC1*** | ***ΔLmxPKAC2*** | ***ΔLmxPKAC3*** |
|---|---|---|---|---|---|
| **amplitude** | 0.0538 | 0.3556 | ***<br><0.001 | *<br>0.0178 | 0.9881 |
| **frequency** | 0.2564 | 0.0551 | ***<br><0.001 | 0.2659 | **<br>0.0035 |
| **w/f** | 0.9664 | 0.2048 | 0.5357 | *<br>0.0093 | 0.3456 |
| **F-length** | 0.4636 | 0.7211 | 0.4434 | 0.1858 | *<br>0.0089 |
