## Supplementary_Figures_S1-S9_and_Supplementary_Data_Legends for "Divergent Protein Kinase A contributes to the regulation of flagellar waveforms in *Leishmania mexicana*"

Fochler et al., 2025.

**Supplementary Figures**

**Supplementary Figure 1. PKAC and PKAR gene synteny.** Cartoon representation of genomic regions for **(A)** PKAC1 and PKAC2, **(B)** PKAC3, **(C)** PKAR1 and **(D)** PKAR3 in representative kinetoplastida species. Gene annotations taken from TritrypDB release 56. The red labels "PKAC1" or "PKAC2" indicate where the names of isoforms differ from the gene names of *L. mexicana*. The colour coding represents the open reading frames (ORF) of syntenic genes within each genomic locus. Green shading highlights genes encoding our PKA subunits of interest. Genomic loci are not to scale. Blunt end and pointed end of coloured bars represent the transcription start and end of the ORFs respectively.

**Supplementary Figure 2. PKAC alignments.** Cartoon representation of the classical protein architecture of *L. mexicana* PKAC subunits compared to the *Homo sapiens* reciprocal best blasts (RBB) of *Lmx*PKAC1 and *Lmx*PKAC3 (accessions NP_001362493 and NP_005035 respectively). Protein domains are colour coded and information was sourced by using InterProScan. Binding site: IPR017441, active site: IPR008271, protein kinase domain: IPR000719, AGC-kinase: IPR000961. Protein sequence alignments of the **(B)** N-terminus extension, **(C)** binding domain, and **(D)** catalytic core across representative kinetospastid species. **(E)** C-terminus extension of PKAC3 homologues in comparison to *Lmx*PKAC1 and -C2 sequences. Underlined sequences in green indicated conservation across PKAC1 and -C2 in all representative species. Underlined residues in yellow indicates conservation across all PKAC3 sequences. Underlined residues in light blue indicates the FXXF feature in the AGC-kinase family (Lilian C. Etchebehere et al. 1997).

**Supplementary Figure 3. PKAC1, PKAC2 and PKAC3 localisation through the promastigote cell division cycle.** Promastigotes in different stages of the cell cycle, expressing **(A)** mNG::PKAC1, **(B)** mNG:: PKAC2 and **(C)** PKAC3::mNG. The first column displays merged micrographs of fluorescence of Hoeschst DNA stain (magenta), mNG (green) and phase contrast, centre column shows the mNG fluorescent signal pseudo-coloured in grey with DNA stain and the last column displays mNG fluorescent signal pseudo-coloured in grey. F, flagella; N, nuclei; N*, dividing nucleus, K, kinetoplasts, nol, nucleolus.

**Supplementary Figure 4. Localisations of *Leishmania* PKAC and PKAR subunits tagged at opposite end.** Fluorescent signal imaged in live promastigote cells expressing **(A)** PKAC1::mNG (LmxM.34.4010), **(B)** PKAC2::mNG (LmxM.34.3960), **(C)** PKAC3::mNG (LmxM.18.1080), **(D)** mNG::PKAR1 (LmxM.13.0160), **(E)** mNG::PKAR3 (LmxM.33.2820).

**Supplementary Figure 5. Localisation of PKAR1 flagella lacking radial spoke and PFR proteins. (A-C)** Micrographs showing the localization of PKAR1::mNG in cells that lack radial spoke proteins: **(A)** Δ*RSP11* (*LmxM.09.1530*) **(B)** Δ*RSP3* (*LmxM.27.0520*) and **(C)** Δ*RSP9* (*LmxM.07.0930*). **(D)** Results of diagnostic PCRs to test for presence of the indicated genes in the parental PKAR1::mNG gDNA and loss of the respective RSP gene in the mutant cell lines. Primers amplifying *PF16* (*LmxM.20.1400*) were used as a positive control for presence of gDNA. **(E-G)** Micrographs showing the localization of PKAR1::mNG in cells that lack parts of the PFR due to deletion of the following genes: **(E)** Δ*PFR2* array (*LmxM.16.1430*) , **(F)** Δ*PFC21* (*LmxM.050950*), **(G)** Δ*PFR-AF1* (PFR assembly factor 1, *LmxM.36.5930*). **(H)** Diagnostic PCRs to test for loss of the respective PFR associated gene in the mutant cell lines, as in D.

**Supplementary Figure 6**. **PCR validation of knock out mutants.** For all gel layouts, the first panel represents a quality control of the primer pairs, whereby PCR amplification using ORF primers listed in figure were used on the parental control genomic DNA template. Following this, the alternation of PCR amplification of a fragment within PF16 locus (LmxM.20.1400) was used as positive control and within the ORF of the target gene of deletion for **(A)** *ΔLmxPKAC3*, *ΔLmxPKA*C2, *ΔLmxPKA*C1, **(B)** *ΔLmxPKAR1*, *ΔLmxPKAR3*. **(C)** double deletion mutants (i) *ΔLmxPKAC1/LmxPKAC2* and (ii) *ΔLmxPKAC1/LmxPKAR3* and **(D)** *ΔLmxPKAR1* and *ΔLmxPKAR3* on parental cell lines expressing mNG C-terminally fused PKAC subunits. The diagnostic primer sequences were taken from LeishGEdit for all but *Lmx*PKAC1 and *Lmx*PKAC2, where a bespoke primer design was required due to sequence similarity. All primer sequences and expected PCR amplicon sizes are listed in Supplementary Table 3A.

**Supplementary Figure 7. Growth curve.** Promastigote cell cultures from *ΔLmxPKAR1, ΔLmxPKAR3, ΔLmxPKAC1, ΔLmxPKAC2* and *ΔLmxPKAC3* alongside parental controls were grown in triplicate flasks from densities $1x10^5$ to $1x10^7$ cells/ml. **(A)** Cell culture densities were recorded with the Cell Counter CASY TTT (OMNI life science) in viable cells/ml every 24 hours from 0 hours to 96 hours. **(B)** Alongside every cell count, the cell population pseudo-diametres in µm were recorded and plotted. **(C)** We calculated the doubling time of the triplicate cell cultures for each population from 0 - 24hrs and 24 - 48hrs then took the average, equating to a total of six doubling times for each deletion mutant population. Error bars in all plots represent standard deviation from the mean. **(D)** Quantification of flagellar length for *Lmx*PKAC and *Lmx*PKAR deletion mutants and parental control populations; n > 70 for each population. Mann-Whitney U test comparing mutant flagellar lengths against parental control measurements.

**Supplementary Figure 8. ΔLmxPKAC1 phenotypic rescue.** Average population swim speed of the parental L. mex Cas9 T7 cell line, the *ΔLmxPKAC1* cell line carrying an episomal addback of the *Lmx*PKAC1 open reading frame (+) and the *ΔLmxPKAC1* cell line carrying an empty pTadd vector (-). The episomal addback restored swim speed to the level of the parental line (p = 0.231, unpaired t-test), **(B)** population overview waveform analysis (POWA) comparing the parental cell line, *ΔLmxPKAC1* with the addback (+) and the *ΔLmxPKAC1* with the empty vector (-). **(C)** Swimming tracks derived from the population motility assays of the three cell lines measured in (A); scalebar 20 µm. **(D)** Results of a diagnostic PCR to test for the reintroduction of the PKAC1 open reading frame. The IFT88 gene was amplified as a control for the presence of gDNA.

**Supplementary Figure 9. Waveform parameter from Fourier analysis.** Scatter plots of flagellum length plotted against waves per flagellum for **(A)** *ΔLmxPKAR1,* **(B)** *ΔLmxPKAR3,* **(C)** *ΔLmxPKAC1,* **(D)** *ΔLmxPKAC2,* **(E)** *ΔLmxPKAC3* and **(D)** parental control. Numbers displayed on plots indicate the $R^2$ correlation coefficient. See Supplementary Table 5 for raw data.

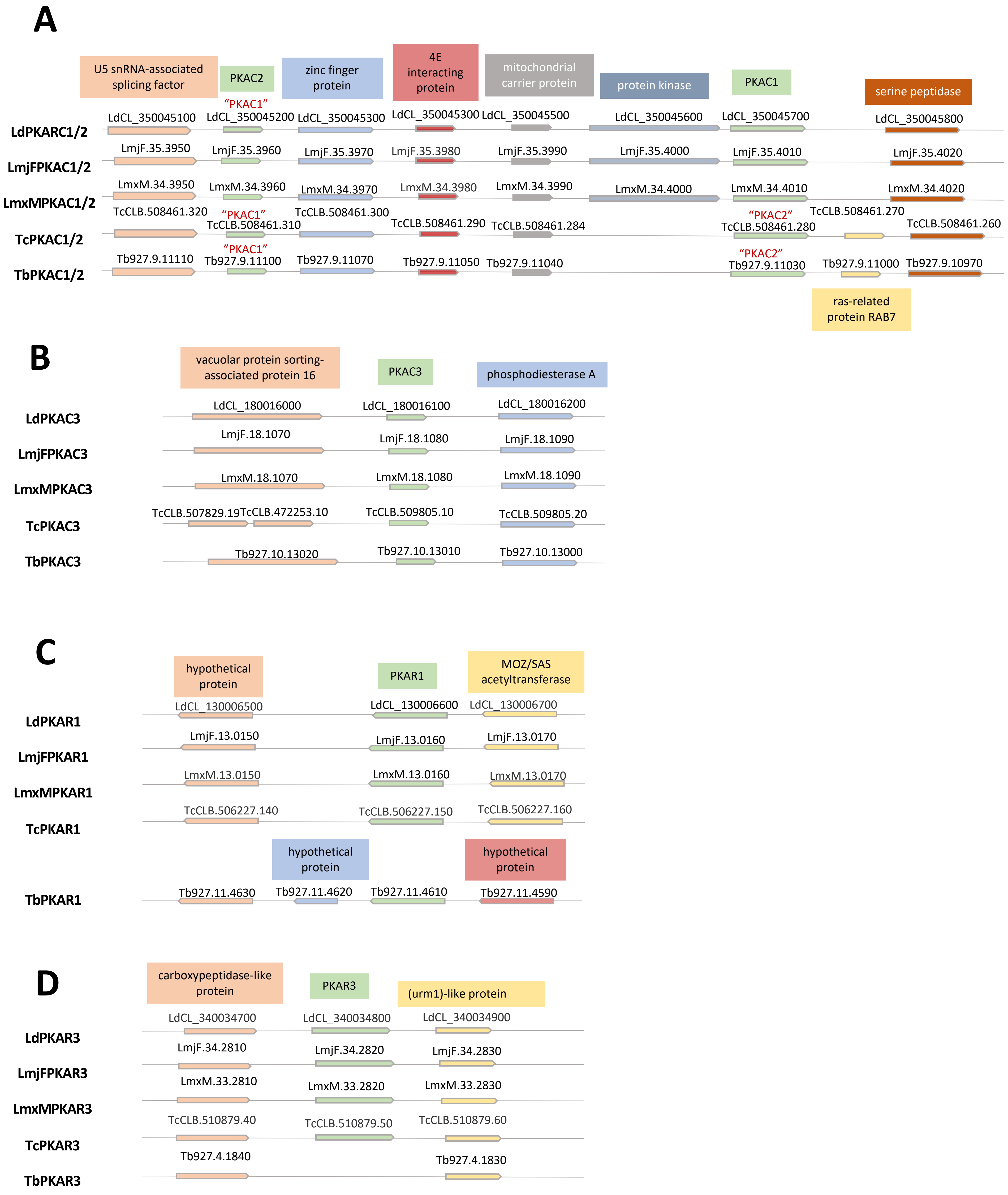
A
U5 snRNA-associated splicing factor
PKAC2
zinc finger protein
4E interacting protein
mitochondrial carrier protein
protein kinase
PKAC1
serine peptidase
LdPKARC1/2
LdCL_350045100
"PKAC1"
LdCL_350045200
LdCL_350045300
LdCL_350045300
LdCL_350045500
LdCL_350045600
LdCL_350045700
LdCL_350045800
LmjFPKAC1/2
LmjF.35.3950
LmjF.35.3960
LmjF.35.3970
LmjF.35.3980
LmjF.35.3990
LmjF.35.4000
LmjF.35.4010
LmjF.35.4020
LmxMPKAC1/2
LmxM.34.3950
LmxM.34.3960
LmxM.34.3970
LmxM.34.3980
LmxM.34.3990
LmxM.34.4000
LmxM.34.4010
LmxM.34.4020
TcPKAC1/2
TcCLB.508461.320
"PKAC1"
TcCLB.508461.310
TcCLB.508461.300
TcCLB.508461.290
TcCLB.508461.284
"PKAC2"
TcCLB.508461.280
TcCLB.508461.270
TcCLB.508461.260
TbPKAC1/2
Tb927.9.11110
"PKAC1"
Tb927.9.11100
Tb927.9.11070
Tb927.9.11050
Tb927.9.11040
"PKAC2"
Tb927.9.11030
Tb927.9.11000
Tb927.9.10970
ras-related protein RAB7
B
vacuolar protein sorting-associated protein 16
PKAC3
phosphodiesterase A
LdPKAC3
LdCL_180016000
LdCL_180016100
LdCL_180016200
LmjFPKAC3
LmjF.18.1070
LmjF.18.1080
LmjF.18.1090
LmxMPKAC3
LmxM.18.1070
LmxM.18.1080
LmxM.18.1090
TcPKAC3
TcCLB.507829.19
TcCLB.472253.10
TcCLB.509805.10
TcCLB.509805.20
TbPKAC3
Tb927.10.13020
Tb927.10.13010
Tb927.10.13000
C
hypothetical protein
PKAR1
MOZ/SAS acetyltransferase
LdPKAR1
LdCL_130006500
LdCL_130006600
LdCL_130006700
LmjFPKAR1
LmjF.13.0150
LmjF.13.0160
LmjF.13.0170
LmxMPKAR1
LmxM.13.0150
LmxM.13.0160
LmxM.13.0170
TcPKAR1
TcCLB.506227.140
TcCLB.506227.150
TcCLB.506227.160
hypothetical protein
hypothetical protein
TbPKAR1
Tb927.11.4630
Tb927.11.4620
Tb927.11.4610
Tb927.11.4590
D
carboxypeptidase-like protein
PKAR3
(urm1)-like protein
LdPKAR3
LdCL_340034700
LdCL_340034800
LdCL_340034900
LmjFPKAR3
LmjF.34.2810
LmjF.34.2820
LmjF.34.2830
LmxMPKAR3
LmxM.33.2810
LmxM.33.2820
LmxM.33.2830
TcPKAR3
TcCLB.510879.40
TcCLB.510879.50
TcCLB.510879.60
TbPKAR3
Tb927.4.1840
Tb927.4.1830

**Figure S1**

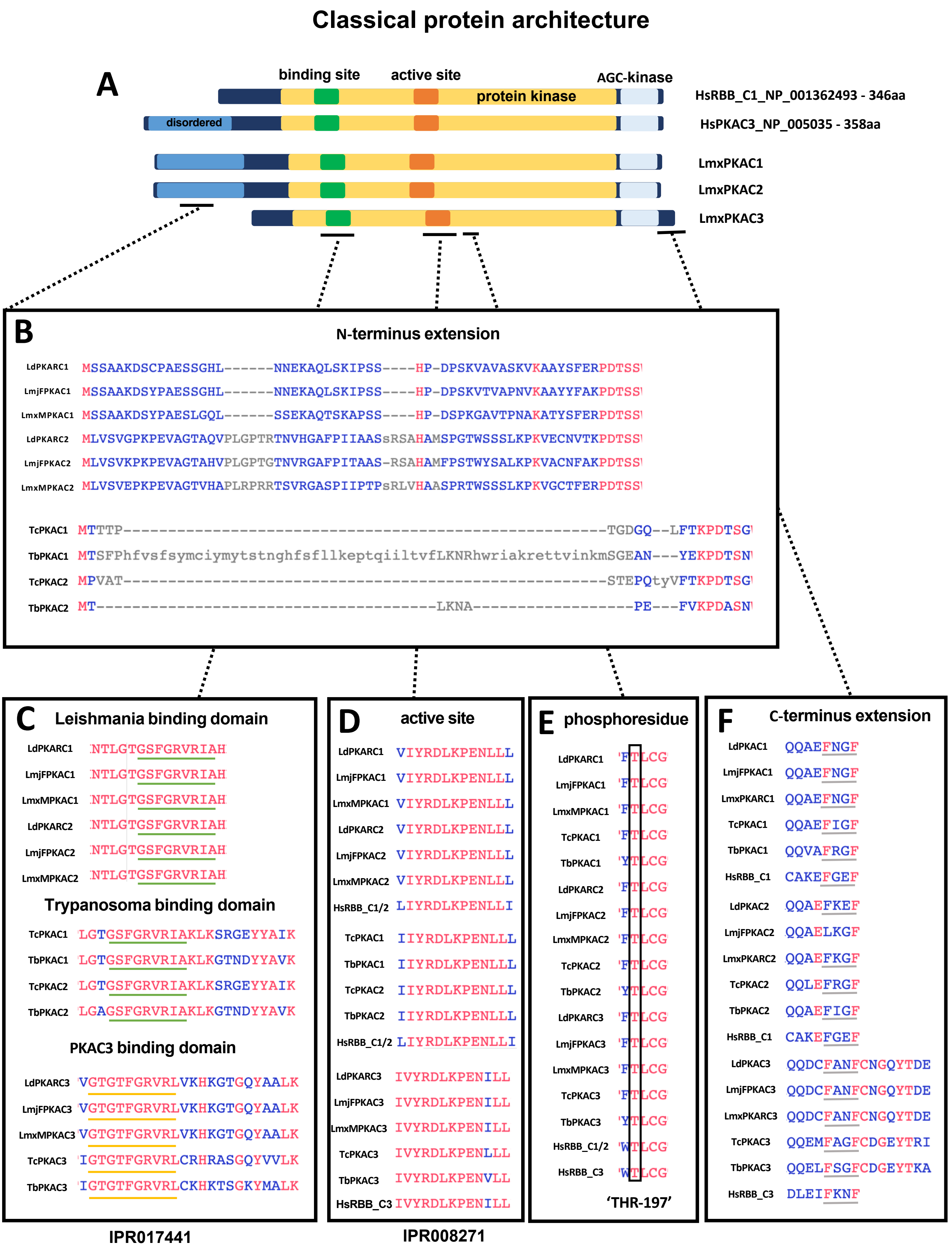

Classical protein architecture
A
binding site
active site
AGC-kinase
protein kinase
HsRBB_C1_NP_001362493 - 346aa
disordered
HsPKAC3_NP_005035 - 358aa
LmxPKAC1
LmxPKAC2
LmxPKAC3
B
N-terminus extension
LdPKARC1 MSSAAKDSCPAESSGHL------NNEKAQLSKIPSS----HP-DPSKVAVASKVKAAYSFERPDTSS
LmjFPKAC1 MSSAAKDSYPAESSGHL------NNEKAQLSKIPSS----HP-DPSKVTVAPNVKAAYYFAKPDTSS
LmxMPKAC1 MSSAAKDSYPAESLGQL------SSEKAQTSKAPSS----HP-DSPKGAVTPNAKATYSFERPDTSS
LdPKARC2 MLVSVGPKPEVAGTAQVPLGPTRTNVHGAFPIIAASsRSAHAMSPGTWSSSLKPKVECNVTKPDTSS
LmjFPKAC2 MLVSVKPKPEVAGTAHVPLGPTGTNVRGAFPITAAS-RSAHAMFPSTWYSALKPKVACNFAKPDTSS
LmxMPKAC2 MLVSVEPKPEVAGTVHAPLRPRRTSVRGASPIIPTPsRLVHAASPRTWSSSLKPKVGCTFERPDTSS
TcPKAC1 MTTTP---------------------------------------------------TGDGQ--LFTKPDTSG
TbPKAC1 MTSFPhfvsfsymciymytstnghfsfllkeptqiiltvfLKNRhwriakrettvinkmSGEAN---YEKPDTSN
TcPKAC2 MPVAT---------------------------------------------------STEPQtyVFTKPDTSG
TbPKAC2 MT--------------------------------------LKNA------------------PE---FVKPDASN
C
Leishmania binding domain
LdPKARC1 NTLGTGSFGRVRIAH
LmjFPKAC1 NTLGTGSFGRVRIAH
LmxMPKAC1 NTLGTGSFGRVRIAH
LdPKARC2 NTLGTGSFGRVRIAH
LmjFPKAC2 NTLGTGSFGRVRIAH
LmxMPKAC2 NTLGTGSFGRVRIAH
Trypanosoma binding domain
TcPKAC1 LGTGSFGRVRIAKLKSRGEYYAIK
TbPKAC1 LGTGSFGRVRIAKLKGTNDYYAVK
TcPKAC2 LGTGSFGRVRIAKLKSRGEYYAIK
TbPKAC2 LGAGSFGRVRIAKLKGTNDYYAVK
PKAC3 binding domain
LdPKARC3 VGTGTFGRVRLVKHKGTGQYAALK
LmjFPKAC3 VGTGTFGRVRLVKHKGTGQYAALK
LmxMPKAC3 VGTGTFGRVRLVKHKGTGQYAALK
TcPKAC3 IGTGTFGRVRLCRHRASGQYVVLK
TbPKAC3 IGTGTFGRVRLCKHKTSGKYMALK
IPR017441
D
active site
LdPKARC1 VIYRDLKPENLLL
LmjFPKAC1 VIYRDLKPENLLL
LmxMPKAC1 VIYRDLKPENLLL
LdPKARC2 VIYRDLKPENLLL
LmjFPKAC2 VIYRDLKPENLLL
LmxMPKAC2 VIYRDLKPENLLL
HsRBB_C1/2 LIYRDLKPENLLI
TcPKAC1 IIYRDLKPENLLL
TbPKAC1 IIYRDLKPENLLL
TcPKAC2 IIYRDLKPENLLL
TbPKAC2 IIYRDLKPENLLL
HsRBB_C1/2 LIYRDLKPENLLI
LdPKAC3 IVYRDLKPENILL
LmjFPKAC3 IVYRDLKPENILL
LmxMPKAC3 IVYRDLKPENILL
TcPKAC3 IVYRDLKPENLLL
TbPKAC3 IVYRDLKPENVLL
HsRBB_C3 IVYRDLKPENILL
IPR008271
E
phosphoresidue
LdPKARC1 FTLCG
LmjFPKAC1 FTLCG
LmxMPKAC1 FTLCG
TcPKAC1 FTLCG
TbPKAC1 YTLCG
LdPKARC2 FTLCG
LmjFPKAC2 FTLCG
LmxMPKAC2 FTLCG
TcPKAC2 FTLCG
TbPKAC2 YTLCG
LdPKARC3 FTLCG
LmjFPKAC3 FTLCG
LmxMPKAC3 FTLCG
TcPKAC3 FTLCG
TbPKAC3 YTLCG
HsRBB_C1/2 WTLCG
HsRBB_C3 WTLCG
'THR-197'
F
C-terminus extension
LdPKAC1 QQAEFNGF
LmjFPKAC1 QQAEFNGF
LmxPKARC1 QQAEFNGF
TcPKAC1 QQAEFIGF
TbPKAC1 QQVAFRGF
HsRBB_C1 CAKEFGEF
LdPKAC2 QQAEFKEF
LmjFPKAC2 QQAELKGF
LmxPKARC2 QQAEFKGF
TcPKAC2 QQLEFRGF
TbPKAC2 QQAEFIGF
HsRBB_C1 CAKEFGEF
LdPKAC3 QQDCFANFCNGQYTDE
LmjFPKAC3 QQDCFANFCNGQYTDE
LmxPKARC3 QQDCFANFCNGQYTDE
TcPKAC3 QQEMFAGFCDGEYTRI
TbPKAC3 QQELFSGFCDGEYTKA
HsRBB_C3 DLEIFKNF


Figure S2

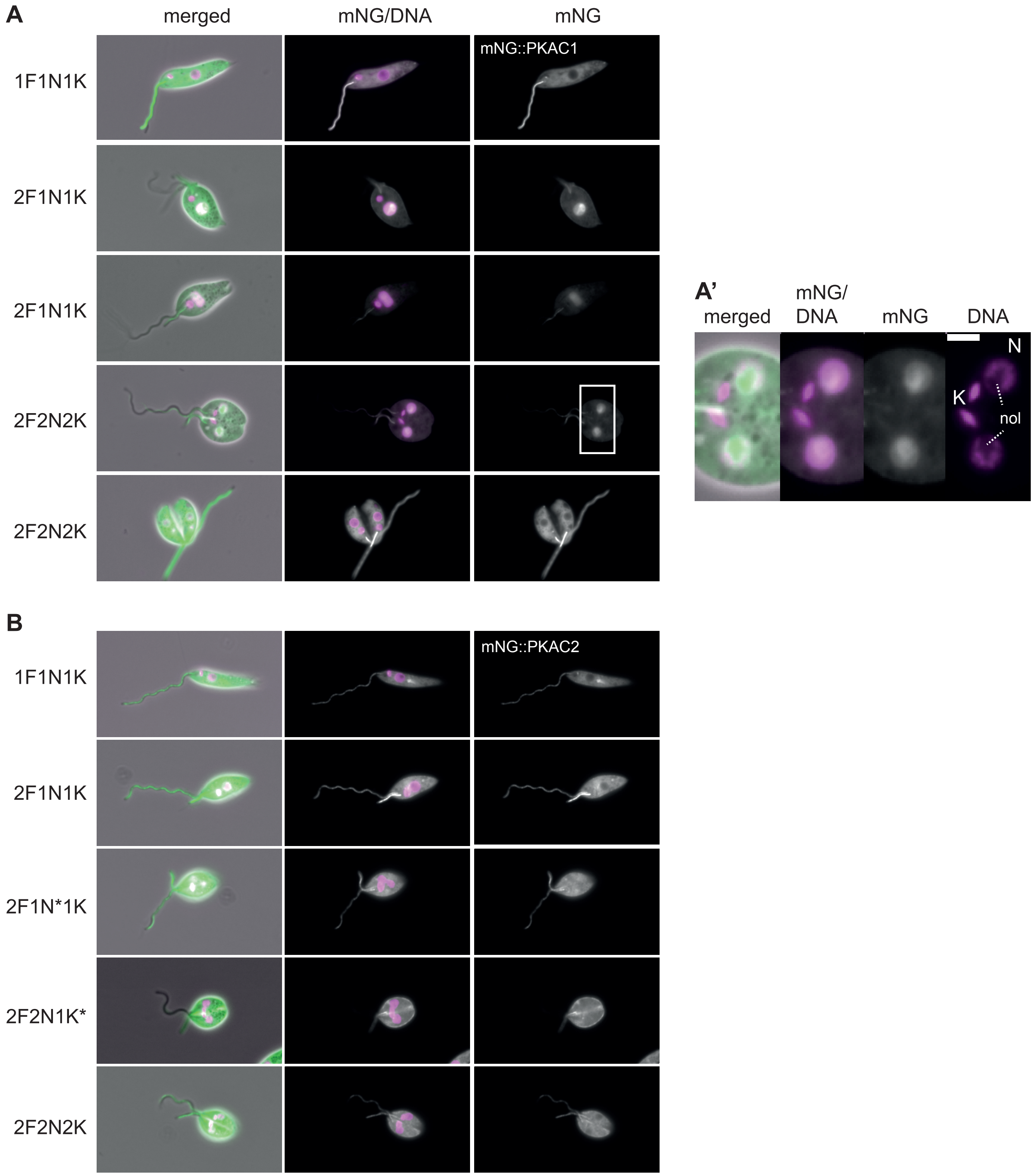
A
merged
mNG/DNA
mNG
mNG::PKAC1
1F1N1K
2F1N1K
2F1N1K
2F2N2K
2F2N2K
A'
merged
mNG/DNA
mNG
DNA
N
K
nol
B
mNG::PKAC2
1F1N1K
2F1N1K
2F1N*1K
2F2N1K*
2F2N2K

**Figure S3**

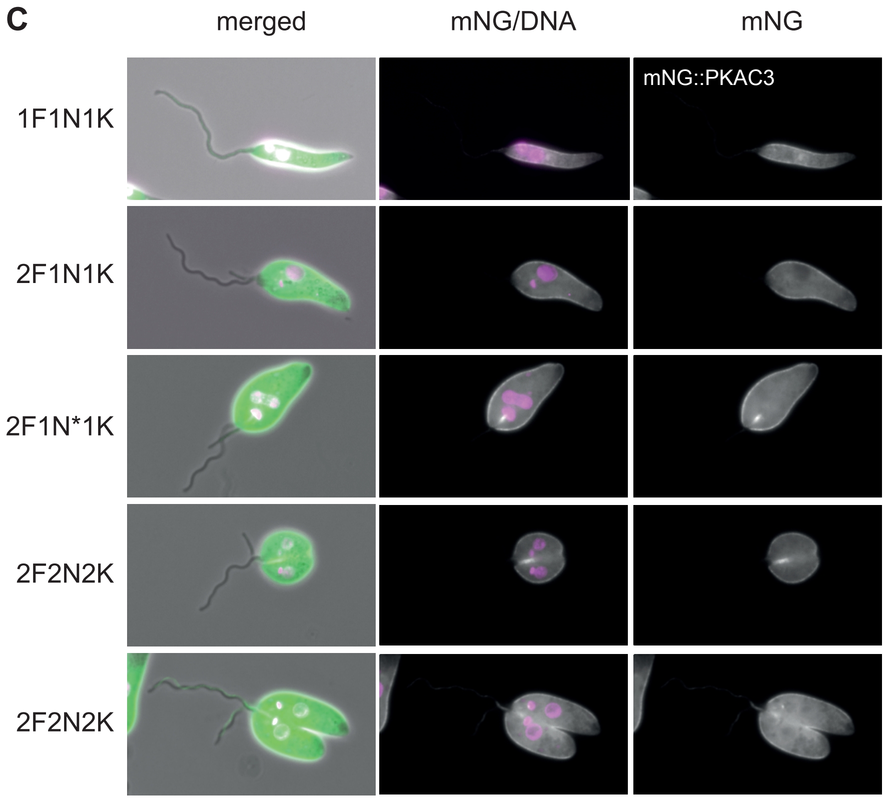
C
merged
mNG/DNA
mNG
mNG::PKAC3
1F1N1K
2F1N1K
2F1N*1K
2F2N2K
2F2N2K

**Figure S3**

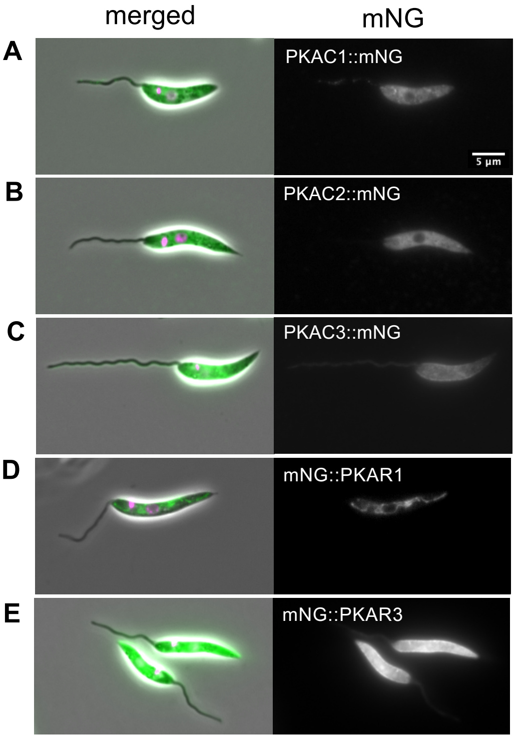
merged
mNG
A
PKAC1::mNG
5 µm
B
PKAC2::mNG
C
PKAC3::mNG
D
mNG::PKAR1
E
mNG::PKAR3

**Figure S4**

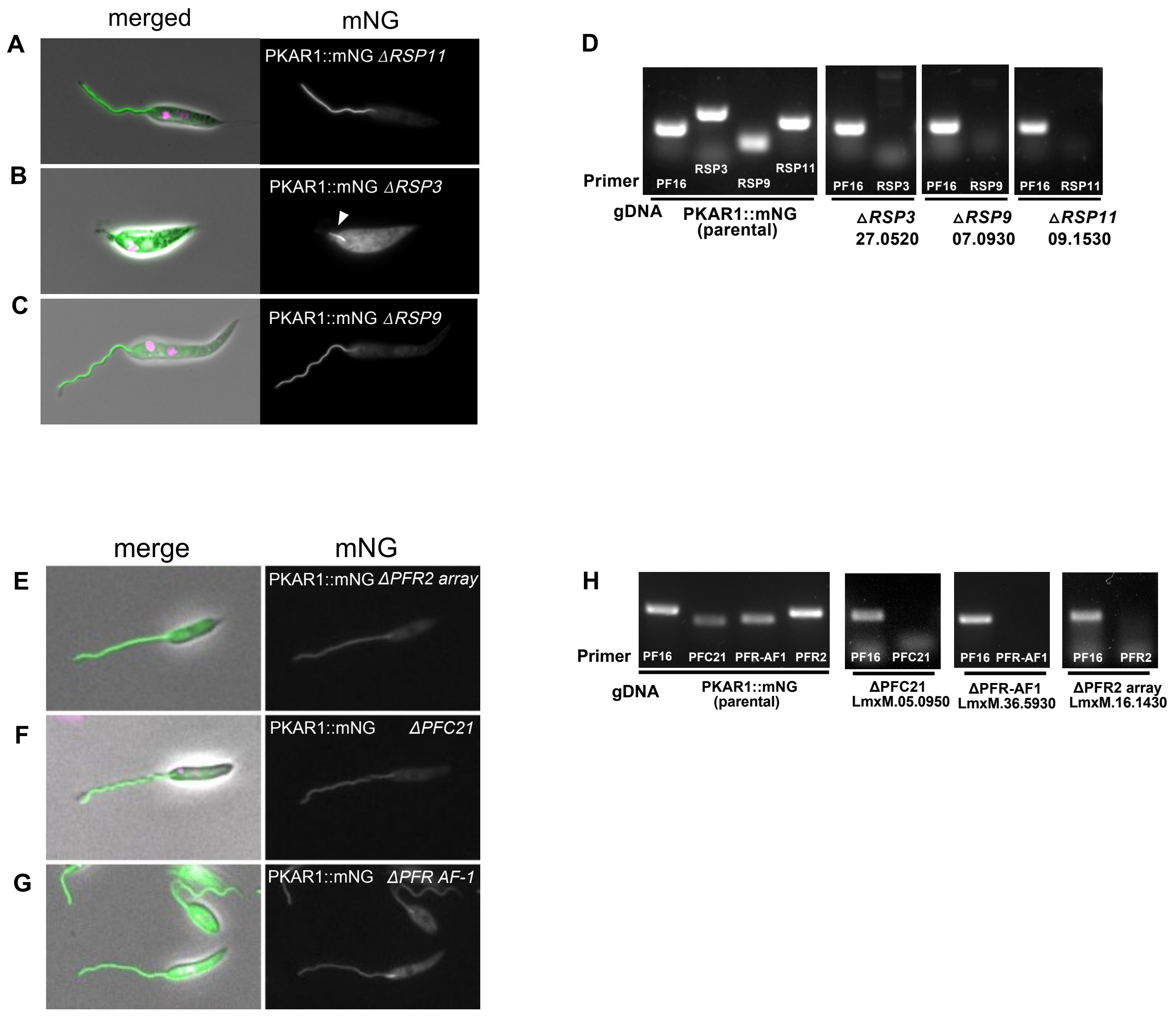
merged
mNG
A
PKAR1::mNG ΔRSP11
B
PKAR1::mNG ΔRSP3
C
PKAR1::mNG ΔRSP9
D
Primer
PF16
RSP3
RSP9
RSP11
PF16
RSP3
PF16
RSP9
PF16
RSP11
gDNA
PKAR1::mNG (parental)
ΔRSP3 27.0520
ΔRSP9 07.0930
ΔRSP11 09.1530
merge
mNG
E
PKAR1::mNG ΔPFR2 array
F
PKAR1::mNG ΔPFC21
G
PKAR1::mNG ΔPFR AF-1
H
Primer
PF16
PFC21
PFR-AF1
PFR2
PF16
PFC21
PF16
PFR-AF1
PF16
PFR2
gDNA
PKAR1::mNG (parental)
ΔPFC21 LmxM.05.0950
ΔPFR-AF1 LmxM.36.5930
ΔPFR2 array LmxM.16.1430

**Figure S5**

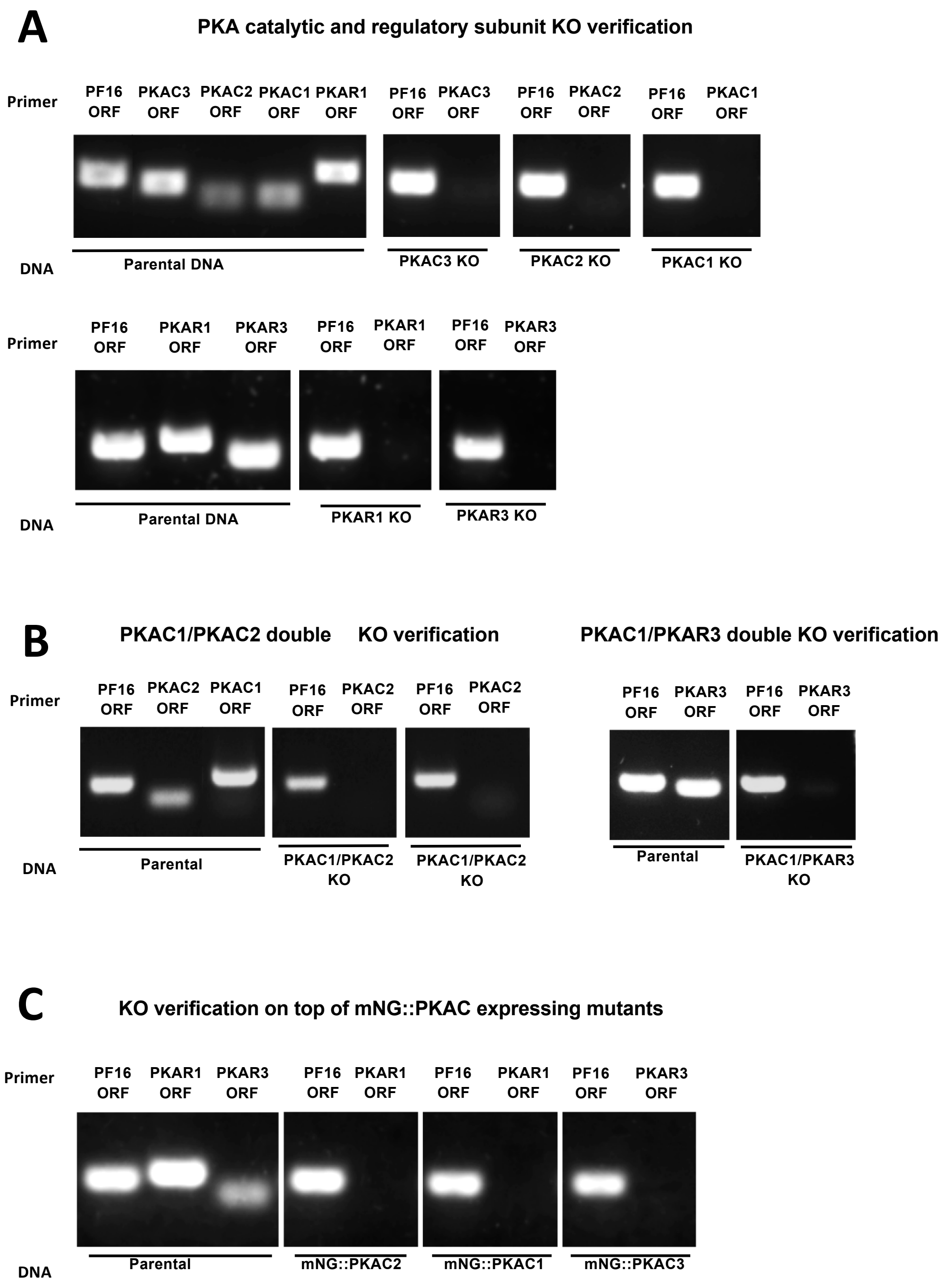


Figure S6

Figure S5

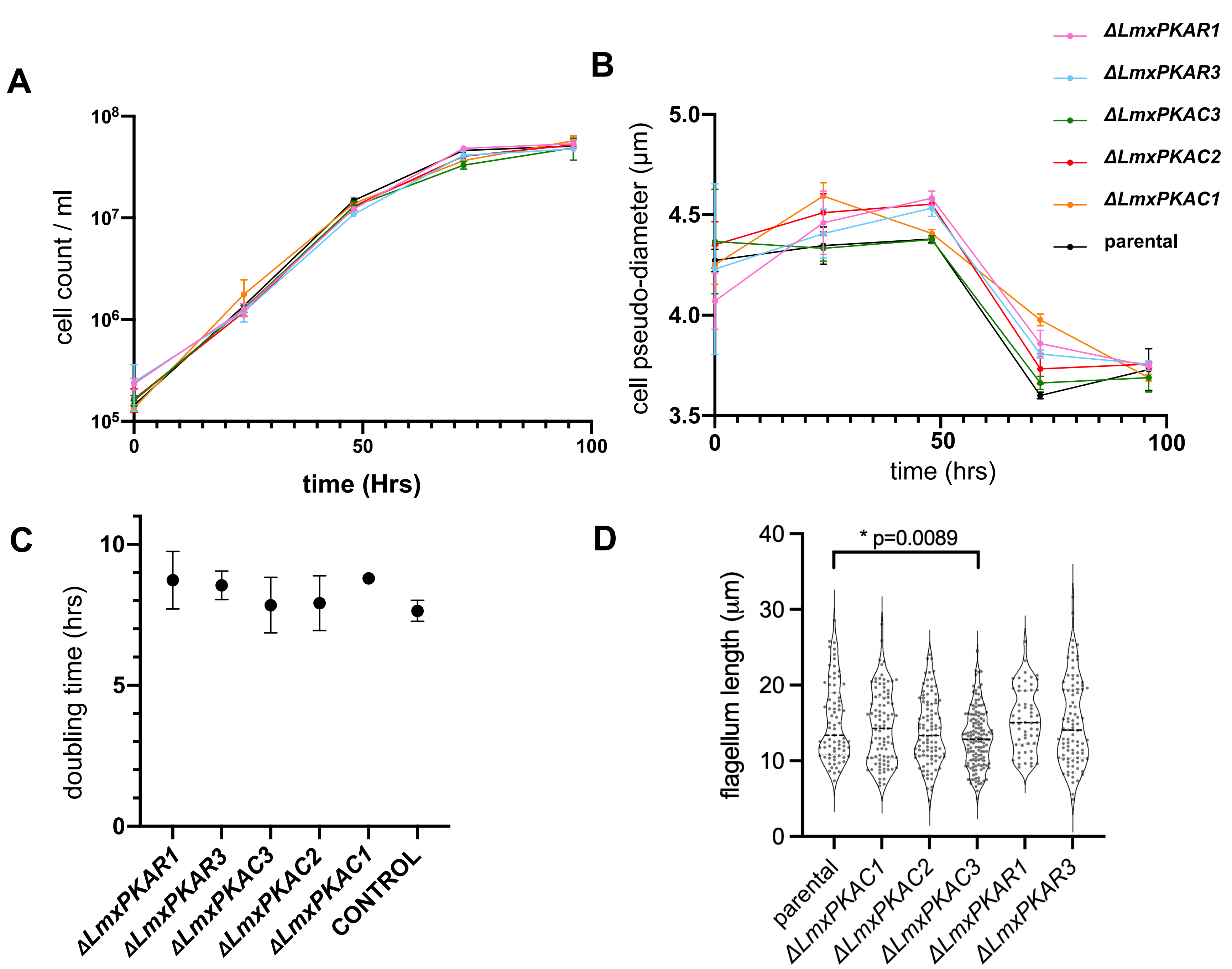
A
cell count / ml
$10^8$
$10^7$
$10^6$
$10^5$
0
50
100
time (Hrs)
B
cell pseudo-diameter (μm)
5.0
4.5
4.0
3.5
0
50
100
time (hrs)
ΔLmxPKAR1
ΔLmxPKAR3
ΔLmxPKAC3
ΔLmxPKAC2
ΔLmxPKAC1
parental
C
doubling time (hrs)
10
5
0
ΔLmxPKAR1
ΔLmxPKAR3
ΔLmxPKAC3
ΔLmxPKAC2
ΔLmxPKAC1
CONTROL
D
flagellum length (μm)
40
30
20
10
0
* p=0.0089
parental
ΔLmxPKAC1
ΔLmxPKAC2
ΔLmxPKAC3
ΔLmxPKAR1
ΔLmxPKAR3

**Figure S7**

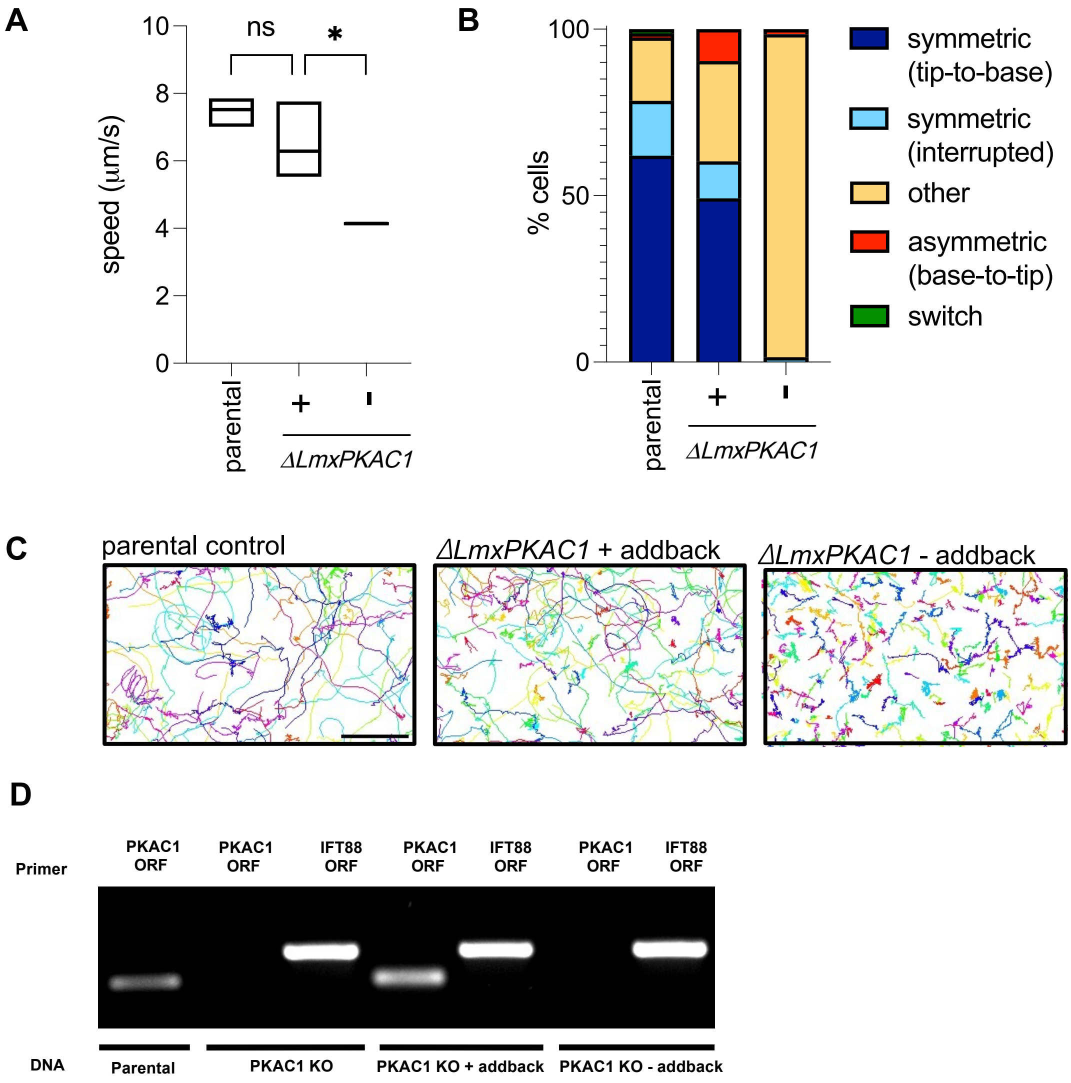
A
10
8
6
4
2
0
speed (μm/s)
ns
*
parental
+
-
ΔLmxPKAC1
B
100
50
0
% cells
parental
+
-
ΔLmxPKAC1
symmetric (tip-to-base)
symmetric (interrupted)
other
asymmetric (base-to-tip)
switch
C
parental control
ΔLmxPKAC1 + addback
ΔLmxPKAC1 - addback
D
Primer
PKAC1 ORF
PKAC1 ORF
IFT88 ORF
PKAC1 ORF
IFT88 ORF
PKAC1 ORF
IFT88 ORF
DNA
Parental
PKAC1 KO
PKAC1 KO + addback
PKAC1 KO - addback

**Figure S8**

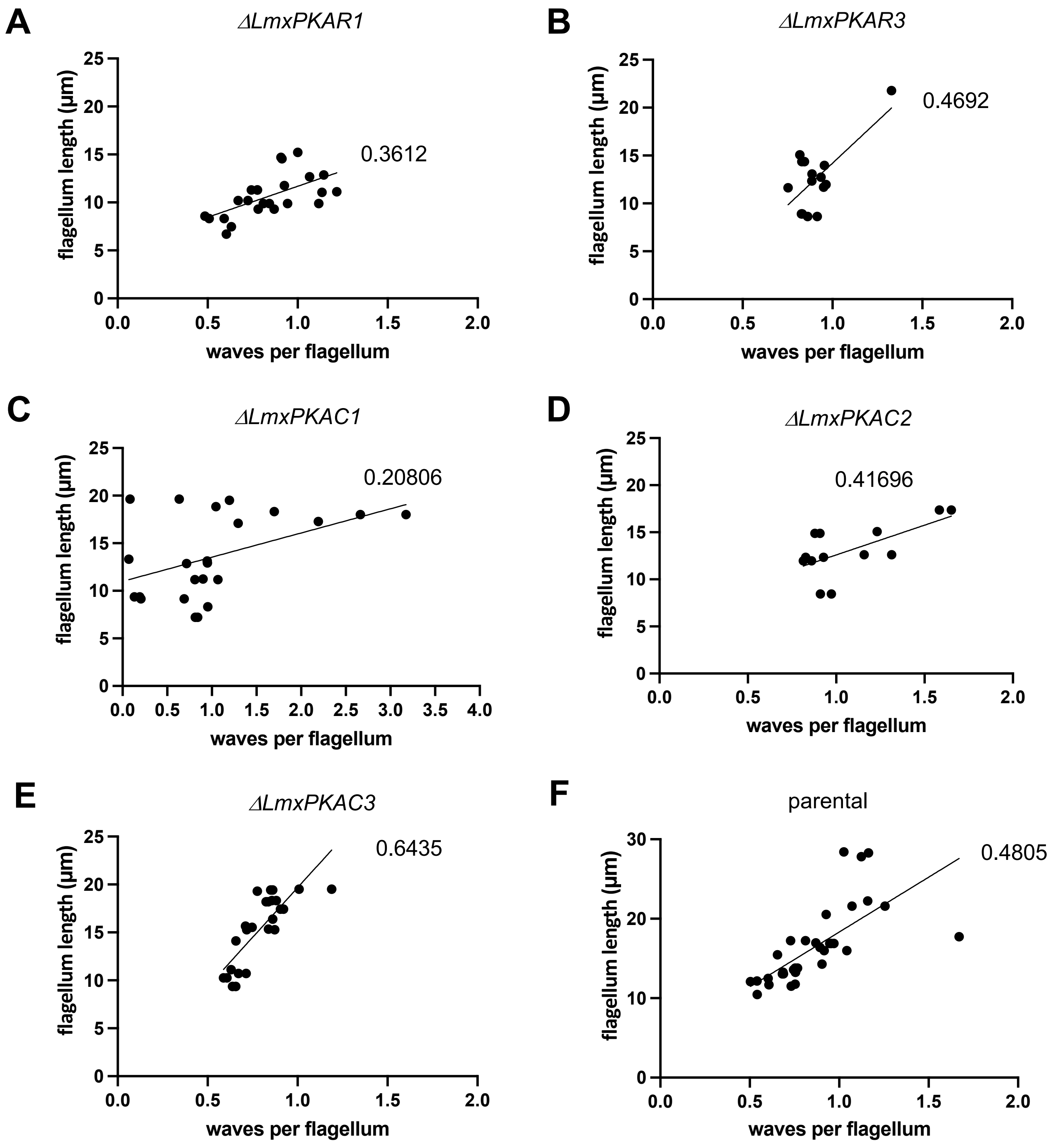
A
ΔLmxPKAR1
0.3612
B
ΔLmxPKAR3
0.4692
C
ΔLmxPKAC1
0.20806
D
ΔLmxPKAC2
0.41696
E
ΔLmxPKAC3
0.6435
F
parental
0.4805
flagellum length (μm)
waves per flagellum

**Figure S9**

**Supplementary Table 1. Population motility raw data.** Raw data for population motility measurements shown in Figure 6A and Supplementary Figure 8A.

**Supplementary Table 2. Waveform parameter statistical test.**

Statistical tests for waveform parameter analysis shown in Figure 7

**Supplementary Table 3. Primers and antibodies.**

Primer sequences for gene deletion validation and details of antibodies used for expansion microscopy.

**Supplementary Table 4. Pharmacological displacement of *Lmx*PKAC1 signal, raw data.**

Raw data for mNG::PKAC1 fluorescence quantification from cytoskeletons treated with different compounds.

**Supplementary Table 5. Waveform parameters, raw data.**

Raw data waveform parameters shown in Figure 7 and Supplementary Figure 9.

**Supplementary movie 1. High-speed videos of parental cells**
Movies of the parental control cell line, captured on a phase contrast microscope at 100 frames per second for a total of 1 second, with one quarter playback speed.

**Supplementary movie 2. High-speed videos of ΔLmxPKAC cells**
Movies of *ΔLmxPKAC1* cells, captured on a phase contrast microscope at 140 frames per second for a total of 1 second, with one quarter playback speed.
